# An 80-channel receive array for 10.5T neuroimaging: Key considerations for SNR optimization

**DOI:** 10.64898/2026.05.06.722982

**Authors:** Matt Waks, Alexander Bratch, Thomas Mercer, Russell Lagore, Steen Moeller, Jeromy Thotland, Lance DelaBarre, Edward Auerbach, Xiaoping Wu, Luca Vizioli, Essa Yacoub, Kamil Ugurbil, Gregor Adriany, Alireza Sadeghi-Tarakameh

## Abstract

**Purpose:** High-density RF receive arrays are required to realize the inherently available SNR and parallel imaging advantages at ultrahigh field strengths, which are essential for high-resolution functional and anatomical brain MRI. This study aims to systematically assess the impacts of often-overlooked parasitic losses associated with various RF coil components, as these losses can degrade the realized SNR and cause significant deviation from the ultimate intrinsic SNR (uiSNR; the theoretical upper bound of available SNR). In addition, we seek to detail engineering solutions to each of these loss mechanisms in pursuit of achieving a higher fraction of the uiSNR limit.

**Methods:** A 16-channel loop-folded dipole transceiver array was developed for 10.5T human head applications and paired with a fully-updated 64-channel receive-only loop array. The optimization of the receive array considered several factors, including (but not limited to) coil dimensions to accommodate a larger population, the size and number of loops to enhance SNR and parallel imaging performance, and circuit design strategies to minimize parasitic losses. The SNR and parallel imaging performance of the receive array were quantitatively assessed by comparison with the uiSNR, as well as existing high-channel-count receive arrays at 7T and 10.5T. Finally, the complete 16-channel transmit, 80-channel receive coil array was safety validated for human use and employed for high-resolution functional and anatomical MRI at 10.5T.

**Results:** Initial results show that the 80-channel array, featuring larger loops in an overlapped layout with optimized circuitry, significantly improves the SNR and approaches the uiSNR limit in a large fraction of the head, while maintaining or enhancing the parallel imaging performance compared to previously used non-overlap layout.

**Conclusion:** This study suggests that, although the traditionally used high-channel-count loop receive array technology can approach the uiSNR limit in the >10T regime, meticulous design optimization—including systematic assessment and minimization of parasitic losses—has become increasingly critical for achieving this goal in this new field-strength territory.

## 1 INTRODUCTION

Magnetic resonance Imaging (MRI) is a powerful, non-invasive imaging modality with excellent soft-tissue contrast. As with any imaging modality, MRI has been subject to incessant efforts to improve its capabilities, which include a drive towards higher spatio-temporal resolutions, and contrast-to-noise ratio (CNR). These efforts in part include the pursuit of higher static magnetic fields (B_0_) in the Ultra-High field (UHF) regime (defined as ≥ 7T), which offer supralinear signal-to-noise ratio (SNR) gains with increasing B_0_ (e.g.,^1–5^), such SNR gains are a fundamental requirement for spatial resolution, scan efficiency, and CNR. Consequently, we witness in recent years a steady increase in the number of UHF MRI systems sited worldwide with further expansion in both the clinical and biomedical research settings expected with the regulatory approval of commercial 7T systems with parallel transmit (pTx) technology (e.g.,^6–8^). Recently, these UHF ambitions have targeted even higher magnetic field strengths in the >10T regime,^9–11^ for additional substantial gains, beyond what 7T has been able to deliver to date.

The benefits of UHF MRI do not come without thier own set of challenges, which can range from patient safety concerns to image uniformity issues (e.g.,^12–18^). In order to overcome these challenges and optimize the use of these powerful systems, more advanced imaging techniques, as well as more sophisticated instrumentation, particularly radiofrequency (RF) coils had been under continuous development (e.g.,^19,20^). Consequently, RF coil technology has undergone an incredible transformation from its roots as singular coil element,^21^ both transmitting RF power for signal excitation and signal reception, to modern large-scale arrays comprising 10’s to 100’s of coil elements (e.g.,^2,3,22–29^) dedicated to signal reception employed together with separate multi-channel transmit or transceiver arrays.^2,3,30–32^ Local multi-channel transmit (Tx) arrays,^33–38^ such as those built for head imaging, avoid exposing larger regions of the subject to transmitter energy, deliver the power to local regions where it is desired, and provide the capability for reducing SAR and improve the uniformity of images through pTx techniques.^39–41^ High density and high-channel-count receive-only (Rx) arrays provide increased sensitivity over large regions of interest by combining data acquired from numerous smaller coils, enhancing the SNR compared to a large single coil element that cover the object of interest. However, achievable SNR gains are not unlimited and are bound by the ultimate intrinsic signal-to-noise ratio (uiSNR).^42^ In generic numerical studies performed for systems up to 9.4T,^43^ 32-channel loop arrays were predicted to capture over 90% of central uiSNR and approximately 80% of uiSNR in intermediate regions of a human brain-size spherical sample across different field strengths.

In our recent works on RF coil development for 10.5T neuroimaging applications, we detailed both Tx and Rx array fabrication techniques, described our safety validation process, and demonstrated the utility of high-channel-count Rx arrays when paired with a 16-channel transceiver (Tx/Rx) array.^2^ However, as is the case with any new experimental setup, this initial work had limitations. We observed that our Rx arrays did not follow earlier numerical predictions;^1^ our 64-channel and 112-channel receive-only head array coils (64Rx and 112Rx respectively) captured ∼53% and ∼66% of central uiSNR, respectively at 10.5T while the same coil layouts performed significantly better at 7 Tesla.^1,2^ In these studies, we utilized the 16-channel transmitter array as a transceiver (Tx/Rx) to help boost the SNR, increasing central SNR performance (defined as iSNR/uiSNR where iSNR is the “intrinsic” experimental SNR of the array independent of image acquisition and relaxation parameters) to 71% and 77% for the 16Tx/80Rx and 16Tx/128Rx array configurations, respectively, but still falling short of the numerical predictions (i.e., > 90%). Even more notably, the SNR performance of these high-channel-count receive arrays dropped rapidly to ∼50% when moving outward from the central to intermediate regions of the sample.

There are well justified reasons for pursuing such high-channel-count arrays for human head imaging at high magnetic fields. The peripheral uiSNR is intrinsically very high and capturing it with increased efficiency requires small RF coils tiled over the head resulting in high-channel-count arrays. Numerical studies have shown that, at the surface of a head-sized spherical phantom, a 32-channel loop array captures less than 10% of the uiSNR at field strengths up to 9.4T.^44^ Increasing the number of such receive channels also improves parallel imaging performance,^45,46^ which is needed particularly for achieving high resolutions within reasonable scan time. However, as the size of individual elements of the array tiled over a head-conformal former decreases, sample noise dominance in these elements also decreases with an accompanying relative increase in coil noise attributed to radiation and circuitry losses, resulting in reduced SNR.^3,26,47^

In this study, we undertook a systematic examination of losses in a 64-channel receive array for 10.5T, investigating fully-overlapped larger individual elements versus the previous non-overlapped 64Rx design of smaller coil elements, and schematic and component-level changes to minimize parasitic coil losses, leading to a solution that diminished the effects of radiative and circuit (resistive) losses, and increased sample noise dominance.^48,49^ This new 64Rx receive-only array was also paired with a newly developed, circumscribing 16-channel Tx/Rx array for human brain applications at 10.5 T. The performance of this combined 16Tx/80Rx array is compared both to theoretical uiSNR as well as to our existing body of work at 10.5T and existing coils at 7T. The new optimized 16Tx/80Rx array had significantly improved SNR performance over the entire volume demonstrating major gains in uiSNR captured in the periphery, the intermediate regions and centrally, reaching ∼90% of iSNR/uiSNR in the center. In addition, the new array had better parallel imaging performance compared to the first-generation 16Tx/80Rx array. Experimental data are presented in a phantom mimicking human head size and electrical properties and in the human head.

## 2 METHODS

### 2.1 RF coils

The new, second-generation 16Tx/80Rx (referred to from here on as 16Tx/80Rx_2_) consisted of two subassemblies: a new 16-channel transceiver array (16Tx/Rx_2_) with each element capable of Tx/Rx functions and a fully-updated 64-channel receive-only array (64Rx_2_).

A coil base was designed to repeatably position the 16Tx/80Rx coil assembly at the same location on the patient table. All coil housings and base components were designed and 3D printed in-house by an additive manufacturing process using a polycarbonate filament (PolyMax™ PC, Polymaker, Changshu, China) on either an X1 Carbon (Bambu Lab, Shenzhen, China) or M2+ (AON3D, Montreal, Canada) printer; no surface finishes were applied to the coil housings prior to the coil assembly process.

#### 2.1.1 Receive-only array

##### Coil loss investigation

Individual RF coil sensitivities can be estimated based on the quality (Q) factor, which can be measured on the electrical bench. The practical losses that limit the Q-factor of each RF coil can be attributed to two main categories: parasitic and sample. In this work, the parasitic electrical losses associated with tuning, matching, detuning, and radiation, as well as the coil fabrication technique and construction materials were investigated. Following the methodology introduced by Chen et al.,^50^ Q-factor measurements were obtained and combined with a hybrid experimental-computational framework to estimate the individual loss components and, consequently, their contributions to the Noise Power Ratio^51^ (NPR) under various conditions, including schematic variations, electrical components, conductor types, RF cables, and other materials, and across multiple coil sizes. Here, 3 × 5 cm^2^ and 5 × 5 cm^2^ loops were selected as control geometries (Figure 1, top row), as these dimensions represent the majority of the coil elements in our first-generation 64-channel Rx array (64Rx_1_), and in the proposed, second-generation 64-channel Rx array (64Rx_2_) detailed below.

**Figure 1:**
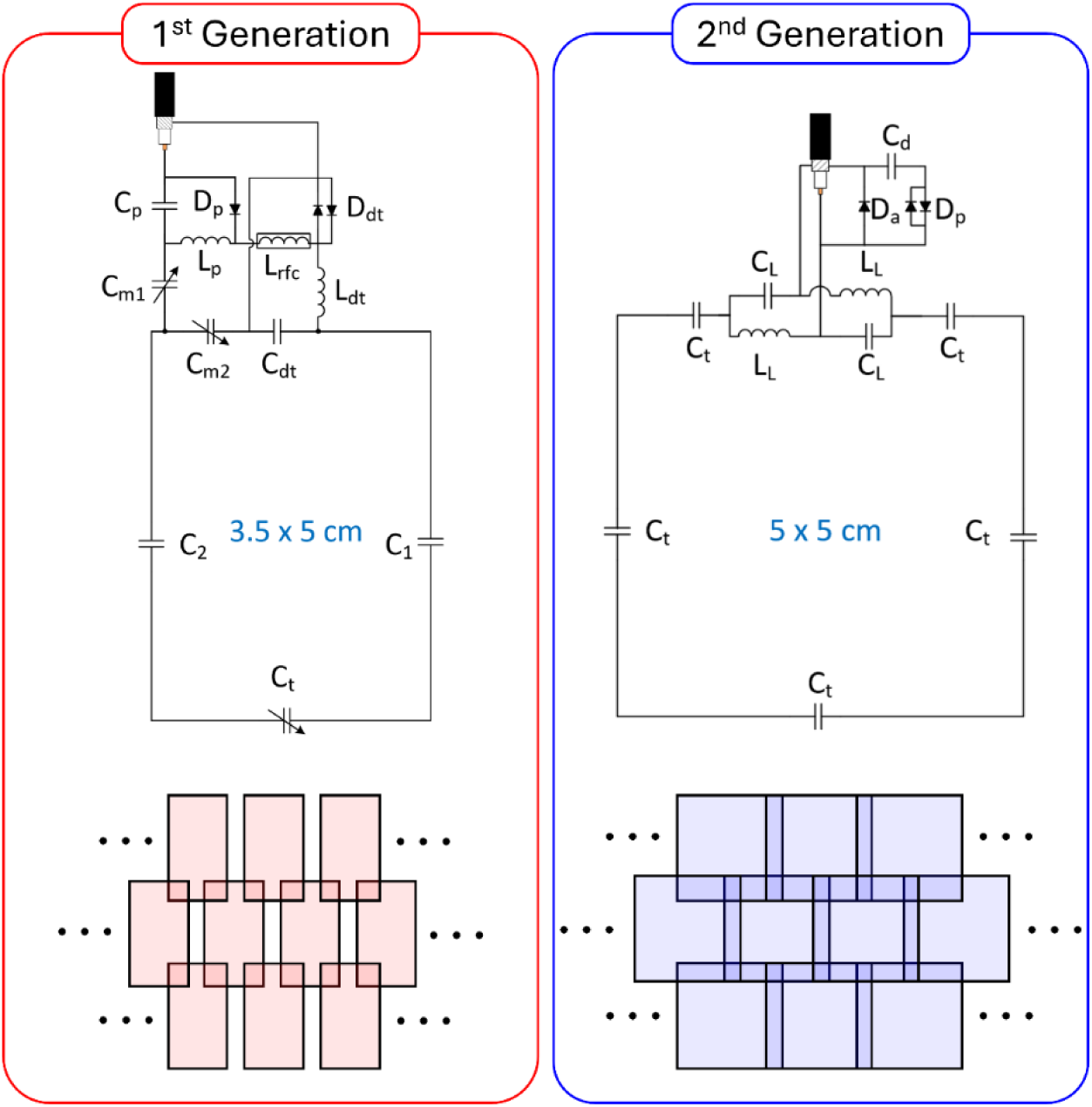
Comparison of two receive-only coil elements, (Left) 1st generation 3x5 cm element from prior work, (Right) 2nd generation 5x5 cm element. (Top) Electrical schematics, (Bottom) element layout of overlapped vs. non-overlapped designs of 64Rx arrays.

Four different components were assumed to contribute to the parasitic losses: conductor loss (a combination of conductor and substrate losses), radiation loss, lossy lumped elements used primarily for coil tuning (mainly capacitors), lossy matching networks (typically comprising capacitors and inductors), and lossy detuning networks (typically comprising a PIN diode and a tank circuit). These losses were estimated for both 3 × 5 cm^2^ and 5 × 5 cm^2^ control loops using a hybrid experimental-computational Q-based technique, as described in detail in.^47^

In brief, the unloaded-to-loaded Q ratio (Q_R_) was measured on the bench for both loops in their resonant state, without matching or detuning networks connected, at various coil-sample distances (1 cm to 4 cm). Electromagnetic modeling of the same configurations was then performed to match the measured Q_R_ values. Knowing the loss associated with the tuning capacitors (*R_cap_* ) used for tunning of the coil, the resistances associated with the sample (*R_Sample_*), conductor (*R_Cond_*), and radiation (*R_Rad_*) were subsequently calculated from the simulations. In the next step, the matching network was added and Q_R_ measurements were repeated, enabling estimation of the loss associated with the matching network (*R_Match_*). Finally, the detuning network was added, and after repeating the Q_R_ measurements, the detuning-network loss (*R_Detune_*) was determined. All technical details of the Q_R_ measurements, as well as the hybrid experimental-computational framework used to quantify parasitic losses, are provided in.^47^

Once all losses were estimated, the Noise Power Ratio (NPR) for each individual loss component was calculated using the following expression to identify its contribution to the thermal noise of the receive coil:

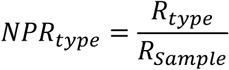

Further details can be found in the Supporting Information.

##### Coil construction

Similar to our previous work,^2,3^ the 64Rx_2_ array (Figures 2, 3D) was designed to fit a human head–conformal former which differed from our previous design by removing the coil elements from over the eyes, and subsequently extending the superior edge to cover more of the frontal cortex without covering the eyes (Figure 2A). The inner surface of this former measured 22.5 cm (anterior–posterior) by 18.25 cm (left–right) with a wall thickness of 2.5 mm. The receive-only array was laid out on the outer surface of the former and consisted of 64 loops in six rows along the z-axis (Figure 2C). The coil elements were geometrically decoupled^22^ by an overlapped arrangement (Figure 1 bottom-right, 2C) in both the superior–inferior direction (between rows) and—in contrast to our first-generation Rx array (64Rx_1_)—azimuthally within each row. Most of the elements were rectangular-shaped centrally (^∼^50 × 50 mm) with smaller rectangular and trapezoidal elements completing the array to fit the organic shape of the former.

**Figure 2:**
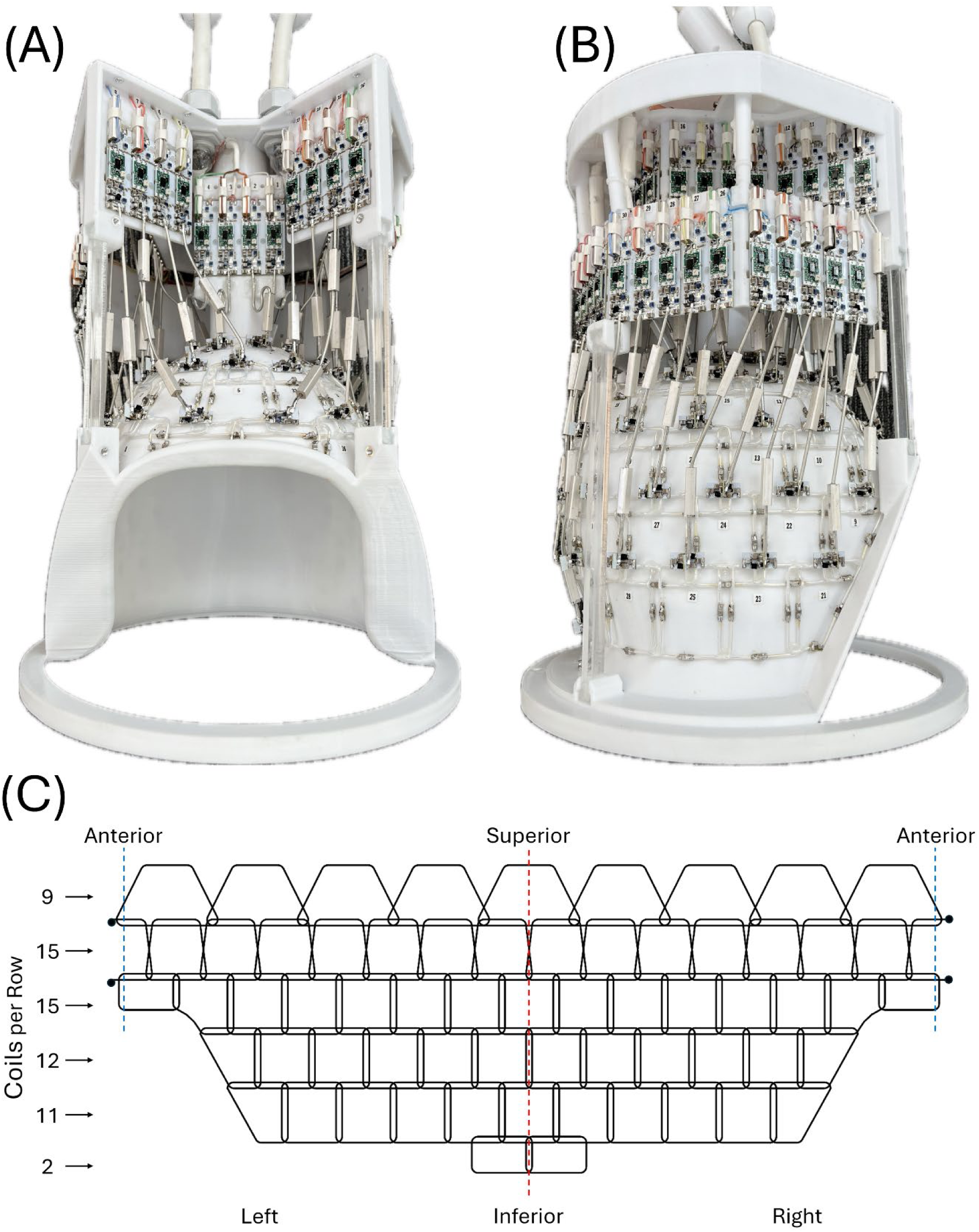
The 2^nd^ generation 10.5T 64-channel receive array (64Rx_2_) as built, viewed from the anterior (A) and patient right (B). (C) Unwrapped 64-channel receive array layout (as viewed from the posterior). The red dashed line represents the posterior, whereas the left and right ends would connect at the anterior (blue dashed line). The number of coil elements per row are indicated on the left.

The 64Rx_2_ coil elements were fabricated using 16 AWG silver-plated copper wire (MWS Wire Industries, Westlake Village, CA, USA) encased in a thin-walled polytetrafluoroethylene (PTFE) tubing (Alpha Wire, Elizabeth, NJ, USA) and formed to fit the organic shape of the former. Each coil element (Figure 1 top-right) was tuned with equally-spaced, evenly-distributed ceramic capacitors (1111 series, Knowles Corp, Itasca, IL, USA), no variable components were used. Passive (UMX9989, Microsemi, Lowell, MA, USA) and active PIN diode (MA4P7464F-1072T, MACOM, Lowell, MA, USA) detuning circuitry was implemented on each element at the lumped element lattice balun^52^ matching circuit. All 64Rx_2_ elements were noise matched to low input impedance preamplifiers (WMAS447A, WanTcom, Chanhassen, MN, USA) via semi-rigid coaxial cables (UT-085C-TP-LL, Micro-Coax Inc., Pottstown, PA, USA) having one or more ceramic quarter-wave cable traps^53^ to reduce interactions within the 64Rx_2_ array as well as with the 16Tx/Rx_2_ array. A preamplifier protection circuit was located at the superior end of the transmission line, prior a phase shifter at the input port of the preamplifier to ensure robustness of the 64Rx_2_ array. All 64 preamplifier circuits were mounted in a serviceable location at the superior end of the 64Rx_2_ array. Both of the 32-channel MR system cable bundles contained multiple cable traps, one electrically-shortened coaxial “bazooka” balun^53^ over the entire bundle, a second over a sub-bundle as the cables were distributed around the assembly, and lastly a ceramic quarter-wave cable trap placed directly at the preamplifier output.

All 64Rx_2_ elements were tuned and matched to 447 MHz (^1^H Larmor frequency at 10.5T) using a 16-port vector network analyzer (ZNBT8; Rhode & Schwarz, Munich, Germany) while loaded with a lightbulb-shaped phantom approximating the size and shape of the head and neck (Figure 3A). The maximum diameter of the phantom in the head region was 17 cm, which tapered to ∼10 cm near the neck, with an overall length (including the neck region) of ∼23 cm. This phantom was filled with a tissue-mimicking polyvinylpyrrolidone (PVP) solution (PVP, 651.1 g/L; NaCl, 17.83 g/L; NiCl2-6H2O, 0.48 g/L; Agar powder, 20 g/L in deionized water).^54^ Using a dielectric probe (DAKS 12, Schmid & Partner Engineering AG, Zurich, Switzerland), both the conductivity and relative permittivity were measured to be 0.65 S/m and 47.2, respectively, at 447 MHz.^1^

**Figure 3:**
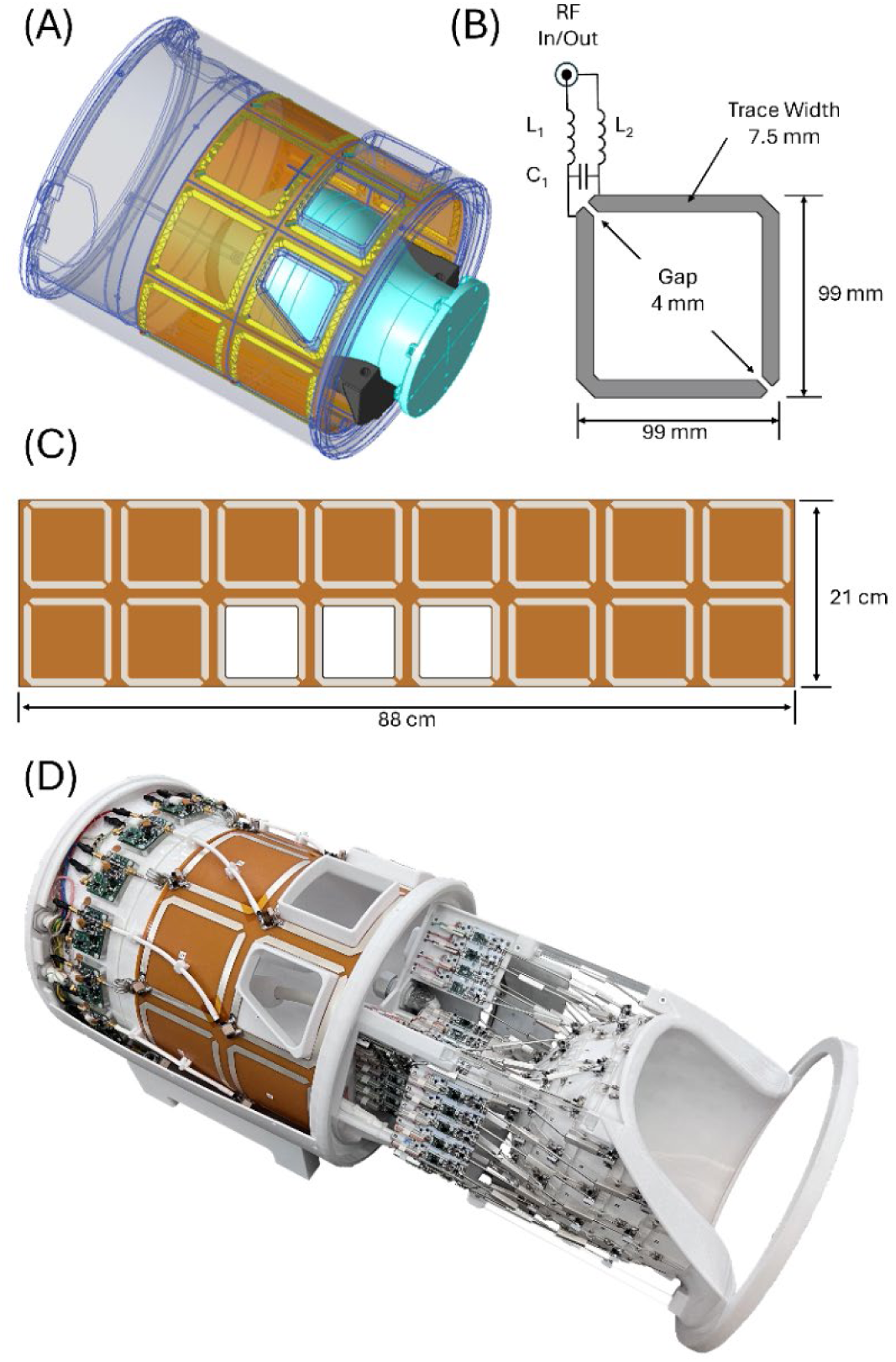
(A) Three-dimensional rendering of the 16-channel transmit with lightbulb phantom & positioner. (B) Schematic of a single loop-folded dipole (LFD) transceiver element. (C) The 16-channel array flex-circuit artwork. (D) The complete 10.5T 16Tx/80Rx_2_ head coil (with cosmetic covers removed) comprising the 16-channel LFD Tx/Rx array containing the integrated system interface and 2^nd^ generation 64Rx_2_ array.

#### 2.1.2 Transceiver array

##### Coil construction

The 16Tx/Rx_2_ array (Figure 3) utilized loop-folded-dipole (LFD) elements, that at 447 MHz, were a direct mechanical replacement to the self-decoupled (SD) elements^55^ detailed in our previous works.^2,3^ This LFD element design further simplified the transmitter electronics beyond the SD elements when compared to traditional loops with distributed capacitors.

Keeping with our previous 16Tx/Rx designs, the transceiver housing (Figure 3A), designed to be compatible with a head gradient, was an open-ended cylinder, 37 cm long with inner and outer diameters of 27.5 and 32 cm, respectively. The 16Tx/Rx_2_ LFD array was laid out in two rows^30,56,57^ (superior-inferior direction) of eight azimuthally distributed elements positioned at a diameter of 28 cm; however, the two rows of the LFD array elements were aligned (Figure 3C) rather than offset by 22.5 degrees (as in previous designs), allowing for inferior element cables to be routed along the virtual ground planes of the superior row of LFD elements (Figure 3D). Similar to the SD design, each element was a 9.15 × 9.15 cm square (conductor center-to-center), with a trace width of 7.5 mm and an 11 mm gap between coil conductors in each row and column. All conductors of the 16-channel SD array were made from 0.5 oz (18 µm) copper with a silver immersion finish on a flexible polyimide substrate with a thickness of 0.1 mm (Figure 3C).

Air core inductors (1812SMS series, Coilcraft, Cary, IL, USA) and Multilayer ceramic chip capacitors (100E Series, American Technical Ceramics, Huntington Station, NY, USA) were used for tuning and matching each LFD element (Figure 3B) at 447 MHz to the lightbulb-shaped phantom described above.

The 16Tx/Rx_2_ LFD array utilized a miniaturized MR system interface integrated into the array housing (Figure 3D), with including Tx/Rx switches with preamplifiers (WMM447P) WanTcom, Chanhassen, MN, USA).^2^ Each of the 16 LFD elements was connected to its respective Tx/Rx switch via ∼17.5 cm of semi-rigid coaxial cable (UT085C-TP-LL, Micro-Coax Inc., Pottstown, PA, USA). In order to mitigate any common mode currents and subsequent coil-to-cable interactions,^58^ the superior element’s cables were wound into a cable trap,^59^ while the inferior row’s cables included a sleeve-type bazooka balun cable trap^53^ and were routed through the virtual ground plane of the anterior elements. The cumulative losses and phase lengths of the complete MR system interface were used to set up the initial B_1_^+^ shim for pTx operation and ensure accurate modeling of our device in EM simulations.

New to this 16Tx/Rx_2_ design was the introduction of openings in the housing in the inferior row of LFD elements over the face (Figure 3). These openings, included with subject comfort in mind, facilitate increased airflow for breathing, and add a feel of openness to reduce the sense of claustrophobia.

##### Safety validation

The 10.5T scanner operates under an Investigational Device Exemption (IDE) from the FDA, requiring each RF coil to be FDA-approved before use with humans. The three-phase EM simulation and safety validation workflow necessary for this process was described in detail in our previous work.^60^ In summary, the exact model of the transmit coil and the uniform light-bulb-shaped phantom was imported into an EM simulation software package (HFSS, ANSYS, Canonsburg, PA). To simplify the simulation, and based on our previous investigations of Tx/Rx coil interactions, the receive coil model was excluded. Initially, all tuning and matching lumped elements were replaced with excitation ports to facilitate the EM/circuit co-simulation technique.^61^ After completion of the EM simulation, the S-matrix was exported to an RF circuit simulation software package (AWR, Cadence, San Jose, CA), where optimization was performed to minimize the differences between the simulated and benchtop-measured S-parameters. After satisfactory agreement was achieved, the optimized lumped-element values were incorporated back into the EM simulation model, and the B_1_^+^ and electric fields were exported for the 16 individual channels of the Tx array. The B_1_^+^ fields were then used to generate the circularly polarized (CP) excitation mode, which was compared with the experimentally measured CP mode. The error between the simulated and measured B_1_^+^ fields was propagated to the predicted error in peak 10 g-averaged local specific absorption rate (pSAR_10*g*_) as described in.^60^ The pSAR_10*g*_ data were calculated from the per-channel electric fields. The estimated pSAR_10*g*_ modeling error (***e_EMM_***) was then combined with intersubject variability (***e_ISV_***) and power monitoring uncertainty (***e_PM_***), as detailed in Steensma, et al.,^62^ to determine a safety factor. This safety factor was ultimately used to scale the virtual observation points (VOPs)^17,63^ to ensure subject safety in compliance with international guidelines^64^.

### 2.2 Experimental characterization

All 10.5T data were acquired on a 10.5 Tesla Siemens MAGNETOM 10.5T Plus; (Siemens Healthineers, Erlangen, Germany) console interfaced to an 88 cm bore 10.5T magnet (Agilent Technologies, Oxford, UK) fitted with a Siemens SC72D gradient coil providing 70 mT/m maximum amplitude and 200 T/m/s slew rate. RF excitation was provided by 16 independent 2-kW RF power amplifiers (Stolberg HF–Technik AG, Stolberg, Germany), and signal reception used 80 of the 128 receive channels developed in-house from Siemens components.

#### 2.2.1 Data for B_1_^+^

Individual complex B_1_^+^ maps for each 16Tx/80Rx array were acquired using the fast relative B_1_^+^ mapping technique (TR = 100 ms, TE = 2.2 ms, flip angle = 15°, voxel size = 2.0 × 2.0 4.0 mm^3^, slices = 15) scaled utilizing the absolute B_1_^+^ maps acquired by actual flip-angle imaging (AFI) technique.^65^ B_1_ mapping parameters of TR_1_ = 25 ms, TR_2_ = 125 ms, TE = 3 ms, flip angle = 60°, voxel size = 1.5 × 1.5 × 5.0 mm^3^, and 100 interleaved axial slices (20% distance factor).

#### 2.2.2 Phantom SNR

Images for SNR comparisons between the first- and second-generation 10.5T 80-channel head coils were obtained with a gradient-recalled echo (GRE) sequence with parameters of TR = 2000 ms, TE = 3.48 ms, nominal FA = 90°, voxel size = 1.0 × 1.0 × 2.0 mm^3^, and 100 interleaved axial slices (20% distance factor) covering the whole phantom. Identical noise images were acquired without an RF excitation pulse and TR = 709 ms.

Flip-angle distribution across the phantom was measured using the AFI technique with the same sequence parameters described in Section 2.2.1.

SNR was calculated following the approach in literature.^2^ To compare the SNR against the uiSNR as well as across field strengths, the measured SNR was converted to the intrinsic SNR (iSNR) in *√HZ/mL* by accounting for sequence parameters, flip angle distribution, and phantom properties, including proton density = 0.69 and T_2_* = 135 ms. For SNR performance calculations (i.e., iSNR/uiSNR), uiSNR was computed using the approach described in.^1^

The calculated iSNR values for both coils were averaged within 0.5-cm-thick concentric shells conforming to the outer contour of the phantom and converging to a 1-cm-diameter spherical volume at the phantom center. This analysis was used to quantify iSNR as a function of depth within the phantom.

#### 2.2.3 In vivo SNR

In vivo data for SNR comparisons were obtained with a gradient-recalled echo (GRE) sequence with parameters of TR = 5000 ms, TE = 3.48 ms, nominal FA = 90°, voxel size = 1.0 × 1.0 × 2.0 mm^3^, and 100 interleaved axial slices (20% distance factor) covering the whole head. Identical noise images were acquired without an RF excitation pulse and TR = 709 ms. Flip-angle distribution was measured using the AFI technique described in Section 2.2.2. Finally, iSNR was calculated by accounting for sequence parameters, flip-angle distribution, and tissue characteristics using the expression below:^2^

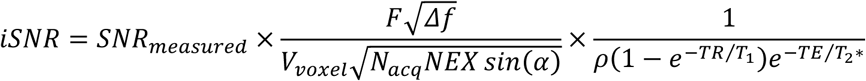

Here, the second term accounts for the sequence parameters and the effect of flip angle, whereas the final term represents tissue-specific parameters, including proton density (*ρ*), T_1_ and T_2_*. At 10.5T, these parameters were measured for gray matter, white matter, and CSF as *ρ* = 0.83/0.70/1.00, T_1_ = 2200/1400/4000 ms, T_2_* = 22.2/18.2/25.0 ms, respectively. To simplify the analysis, a single scaling factor was ultimately used to scale the measured SNR across tissues by averaging the three tissue-specific scaling factors.

#### 2.2.4 g-Factor and parallel imaging

The g-factor maps were calculated from the SENSE equations for simultaneous multislice/multiband (SMS/MB) acceleration with k-space undersampling, and for one-dimensional (1D) and 2D k-space undersampling. The data for the g-factor maps were obtained with a fully sampled 1-mm isotropic 3D Multi-Echo GRE sequence with 8 echoes, TE = 5/15/25/35/45/55/65/75 ms, TR = 80 ms, flip angle = 15°, Pixel Bandwidth = 280 Hz/pixel and 240 slices.

To assess the parallel imaging performance of the first- and second-generation 10.5T 80-channel coils under various undersampling conditions, we evaluated 1D undersampling factors ranging from 2 to 8, as well as SMS/MB acceleration with MB factors of 4 and 8 combined with in-plane undersampling factors of 2 and 4. For visualization, maximum intensity projections (MIPs) of g-factor maps corresponding to SMS/MB data were extracted in the anterior–posterior direction over an 80-mm-thick slab in the sagittal plane and displayed over a silhouette of the central sagittal slice as 1/g maps.

#### 2.2.5 SNR across field strengths

All 7T data were acquired on a 7 Tesla Siemens Terra.X (Siemens Healthineers, Erlangen, Germany) console interfaced to an 88 cm bore 7T magnet (Agilent Technologies, Oxford, UK) fitted with a Siemens SC72D gradient coil providing 135 mT/m maximum amplitude and 250 T/m/s slew rate. RF excitation was provided by 16 independent 2-kW RF power amplifiers (Stolberg HF–Technik AG, Stolberg, Germany), and signal reception used up to 64 receive channels.

Three 7T RF head coils were used to compare SNR across field strengths (7T vs. 10.5T) utilizing the lightbulb-shaped phantom described above, and the same pulse sequence parameters. The first RF coil, currently considered the “industry standard” for 7T, was a commercially available 1Tx/32Rx array (Nova Medical Inc., Wilmington, MA, United States). The second commercially available RF coil, currently considered the “state-of the art” for 7T, was a 16Tx/64Rx array (MR CoilTech Ltd, Glasgow, Scotland). The third RF coil was an in-house custom-built 16Tx/64Rx array designed to schematically and geometrically match the 10.5T 64Rx_1_ as close as possible to compare the sensitivity of the same receiver array design across field strengths; as such, this 64Rx 7T array was not necessarily built to achieve optimal performance at 7T.

SNR data for all 7T coils were acquired as described in Sections 2.2.2 and 2.2.3. The conversion to iSNR was also performed as described in those sections, with the only difference being the phantom and tissue relaxation parameters used for 7T. For the phantom, T_2_* at 7T was extrapolated from the 10.5T data as 202 ms. For in vivo iSNR conversion, the relaxation parameters for gray matter, white matter, and CSF at 7T were obtained from the literature as T_1_ = 1600/1000/3000 ms, T_2_* = 33.3/27.3/37.5 ms, respectively.

For the iSNR-versus-depth analysis, iSNR was averaged within 0.5-cm-thick concentric shells conforming to the outer contour of the head. The final data point on the in vivo iSNR–depth curve was calculated by averaging iSNR within a centrally placed 1-cm-diameter sphere in the cerebrum.

### 2.3 Neuroimaging at 10.5T

All volunteers for this study provided written informed consent. The study was conducted under an Investigational Device Exemption from the FDA and approval from the University of Minnesota IRB, as described previously.

#### 2.3.1 Anatomical imaging

Three different sets of anatomical images were obtained:

a. **Cerebellum-focused GRE:** Cerebellum-focused GRE images were acquired with TR = 600 ms, TE = 20 ms, nominal FA = 40°, in-plane acceleration factor (iPAT) = 2, bandwidth = 40 Hz/pixel, resolution = 0.2 × 0.2 × 1.0 mm^3^, and matrix size = 1024 × 1024. Ten slices were acquired in a sagittal–coronal orientation tilted perpendicular to the tree-like white matter structure of the cerebellum. Data were acquired in 5 minutes and 28 seconds using phase- and magnitude-based RF shimming targeted to produce a uniform transmit field across the cerebellum.
b. **Susceptibility-weighted imaging (SWI):** SWI was performed using a 3D GRE sequence with 36 contiguous axial–coronal oblique slices, 0.21 × 0.21 mm^2^ in-plane resolution, 1.3 mm slice thickness, TR = 35 ms, TE = 18 ms, FOV = 215 × 188 × 46.8 mm, iPAT = 3, nominal FA = 14°, and bandwidth = 315 Hz/pixel. Phase-only RF shimming was performed to target a CP-like transmit field across the whole brain.
c. **T1-weighted whole-brain imaging:** T1-weighted whole-brain images were acquired using an MP2RAGE sequence with 0.7 mm isotropic voxels, TR/TE = 5000/1.97 ms, TA = 7.5 minutes, FOV = 253 × 270 mm^2^, iPAT = 4, inversion times of 840 ms and 2370 ms, and partial Fourier = 6/8. Whole-brain phase- and magnitude-based RF shimming was used for this acquisition.

#### 2.3.2 Functional imaging

Whole-brain resting-state functional MRI (rs-fMRI) data were acquired using a 2D echo-planar imaging (EPI) sequence with 0.6 mm isotropic resolution, corresponding to a voxel volume of 0.216 µL. The sequence parameters were as follows: TR = 3094 ms, TE = 22 ms, echo spacing = 1.24 ms, pixel bandwidth = 930 Hz/pixel, partial Fourier = 5/8, MB = 5, iPAT = 3, and nominal FA = 78°. Whole-brain phase- and magnitude-based RF shimming was used for this acquisition.

Another submillimeter whole-brain resting-state functional MRI (rs-fMRI) dataset, targeting higher temporal resolution, was acquired using a 2D EPI sequence with 0.85 mm isotropic resolution and a volume acquisition time of 1.7 s. The sequence parameters were as follows: TR = 1690 ms, TE = 16.6 ms, echo spacing = 1 ms, pixel bandwidth = 1208 Hz/pixel, partial Fourier = 5/8, MB = 4, iPAT = 4, and nominal FA = 50°. Whole-brain phase- and magnitude-based RF shimming was used for this acquisition.

## 3 RESULTS

### 3.1 Bench Measurements and Safety Validation

#### 3.1.1 Receive-only array

##### Coil loss investigation

The impacts of the various test conditions vs Q-factor are detailed in Supporting Tables S1-S3. Informed by these results, we redesigned the individual Rx coil schematic and compared that to our prior work via the control-sized coils (3 × 5 cm, 5 × 5 cm) (Figure 1), the results can be visualized as contributions to noise power ratio in Figure 4. The data clearly shows the importance of the larger loops—achieved through an overlapped layout—in establishing greater sample noise dominance, whereas in a smaller loop layout at a certain coil-sample distance, the often-overlooked parasitic losses (e.g., losses associated with the coil conductor, distributed capacitors, and preamplifier decoupling circuitry) can become comparable to the sample noise.

**Figure 4:**
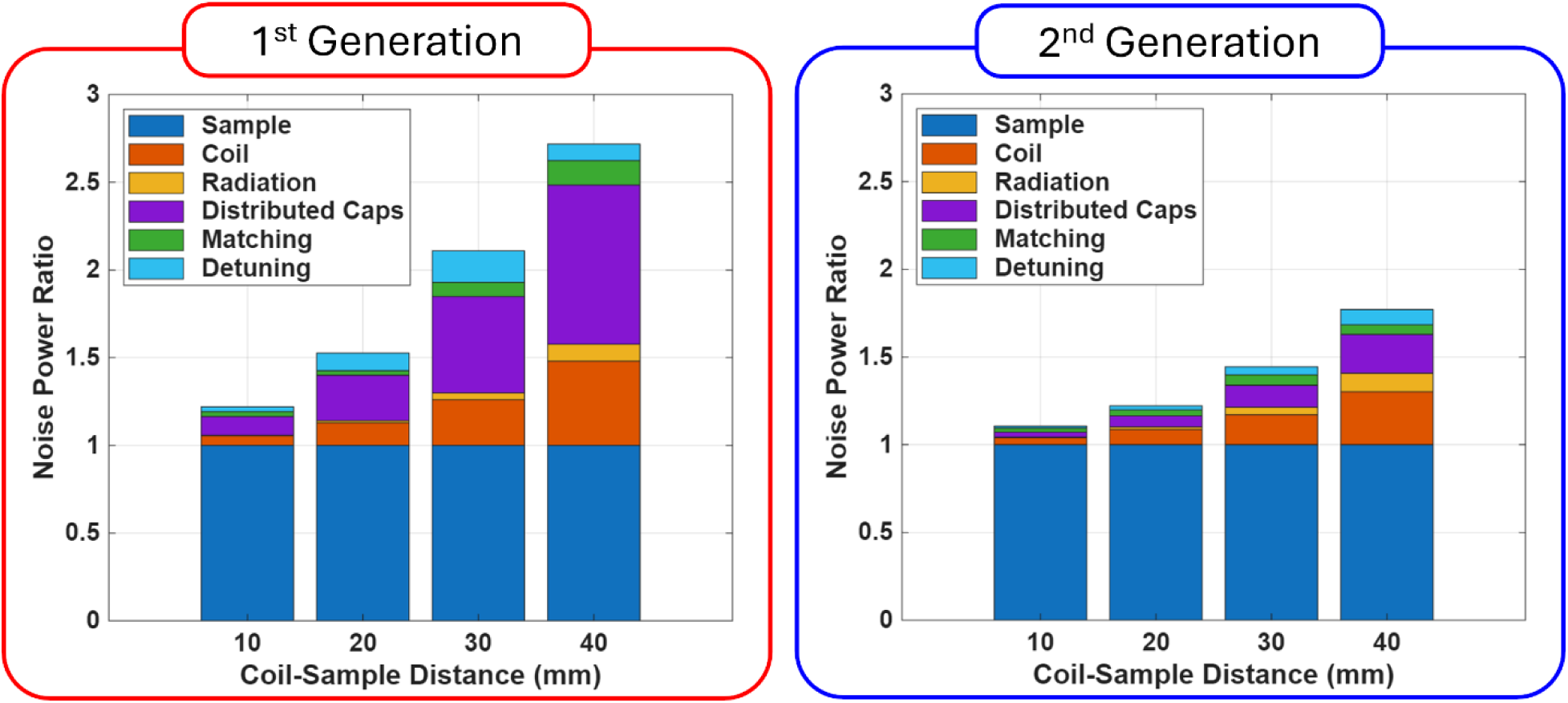
Noise power ratio comparison of two receive-only coil elements, (Left) 1^st^ generation 3x5 cm element from prior work, (Right) 2^nd^ generation 5x5 cm element.

##### 64-channel receive array

All elements in the phantom-loaded 64Rx_2_ array were matched to better than −15 dB with the complete array assembled. It was impractical to measure the complete S-parameter matrix of this 64-element array; however, sample measurements of multiple coils in high density regions of the complete array—considered to be worst case scenario—for overlapped neighboring elements demonstrated a median coupling (S_ij_) of -14.7 dB between rows, and -14.5 dB azimuthally; while the median next-nearest neighbor coupling was observed to be -14.6 between rows, and -19.3 dB azimuthally. This degree of isolation was considered sufficient given that preamplifier decoupling further improves the isolation. Among the multiple-sized array elements, the unloaded-to-loaded Q-ratios (Q_R_) evaluated as Q_unloaded_/Q_Loaded_, as measured with a decoupled probe pair, ranged from 3.7:1 for the smaller lightly loaded coil elements in the inferior row at the posterior to a maximum measured 14.7:1 for a larger heavily loaded element near the superior end of the array. During excitation, a sample 64Rx_2_ element demonstrated active detuning of ∼26 dB, and ∼50 dB of preamplifier protection when combined with the remote diode on the preamplifier circuit assembly.

Each ceramic quarter-wave cable trap on the 0.085” semi-rigid cables between the coil elements are preamplifier provided ∼14 dB of attenuation, while the same ceramic traps placed at the preamplifier output provided ∼25-30 dB attenuation with two turns of the micro-coaxial receive cables. The WMAS447A preamplifiers, which have an input impedance of 1.5Ω, produced a typical gain and noise figure of 28.0 dB and 0.25 dB respectively, and better than -20dB preamplifier decoupling when measured (S_21_) with a decoupled probe pair.

#### 3.1.2 Transceiver array

##### 16-channel transceiver array

Bench measurements demonstrated good tuning and matching for the 16Tx/Rx_2_ array elements for both the phantom load and the human head, with and without the 64Rx_2_ insert. Supporting Figure S2 shows the return losses (coil matching values, S_ii_) for the phantom-loaded 16Tx/Rx_2_ array with the 64Rx_2_ array in place ranged from −16.2 to −38.4 dB, with a median of −18.2 dB. The interelement coupling (S_ij_) ranged from −10.7 to −62.7 dB with a median of −32.3 dB. Most of the coupling within the 16Tx/Rx_2_ array happened between the vertical neighbors, with a median of −11.7 dB; whereas the median coupling between azimuthal and diagonal neighboring elements was -14.3 dB and -20.8 dB respectively.

In the transmit mode, the typical insertion loss of the complete integrated interface including the MR system cables was −0.9 dB with a 0.5° phase balance between the 16 Tx/Rx switches; the preamplifiers were isolated from the transmitter-to-coil path by ∼51.7dB. During acquisition, the WMM447P preamplifiers had a gain of approximately 28.0 dB, noise figure of 0.45 dB, typical insertion losses in the receive path preceding the preamplifier measured −0.4 dB including the semi-rigid coaxial cable connecting the LFD element to the Tx/Rx switch; the transmitter input port was isolated from the receive path by 51.9 dB during acquisition.

##### Safety validation

Figure S4A shows a per-channel comparison of B_1_^+^ between simulation and experiment. Figure S4B shows the measured and simulated B_1_^+^ maps for the CP excitation of the 16-channel LFD array (16Tx/Rx_2_). The normalized root mean square error (NRMSE) between the simulated and measured maps was 35%, and this value was propagated into the corresponding pSAR_10*g*_ error regions, as shown in Figure S4C. The 99.9^th^ percentile of the resulting error distribution is highlighted in the histogram and was defined as *e_EMM_* for the coil. The resulting *e_EMM_* of 46% was then combined, in a sum-of-squares manner, with *e_ISV_* = 50% (from Le Garrec et al.)^66^ and *e_PM_* = 15% (reported by the vendor), yielding an overall safety factor of 1.7.

### 3.2 SNR Characterization

#### 3.2.1 Phantom B_1_^+^

Supporting Figure S3 shows phantom-loaded B_1_^+^ efficiency maps for both the first- and second-generation 16Tx/80Rx coil assemblies. One can observe the higher peak B_1_^+^ per watt within the phantom with the updated array, 16Tx/80Rx_2_. The original 16Tx/80Rx_1_ array produced B_1_^+^ efficiency values of 0.313 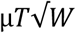 while the 16Tx/80Rx assembly produced 0.414 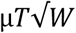 averaged across a 1 cm spherical volume located at the crosshairs

#### 3.2.2 Phantom SNR

Figure S5 shows the experimentally measured noise correlation matrices for the 16Tx/80Rx_2_ array while loaded with the lightbulb-shaped phantom. The average noise correlation was 6.1% with an observed maximum of 51%.

Figure S6A shows SNR maps within the lightbulb-shaped phantom for both the first- and second-generation 64Rx and 80Rx arrays in a central axial slice, demonstrating gains throughout the phantom from utilizing the 16Tx/Rx array during acquisition. The SNR was also plotted as a function of depth (Figure S6C) with SNR averaged in 0.5-cm-thick concentric shells, around a 1-cm-diameter central volume located at the center of the phantom. The additional 16 receive channels from the transceiver array provided an 11% increase in central SNR, as measured within the 1 cm-diameter volume.

The SNR maps from both 16Tx/80Rx arrays are shown in all three orthogonal planes (Figure 5A). The ratio of SNR as a function of depth plotted for SNR averaged in 0.5-cm-thick concentric shells, with a 1-cm-diameter central volume (Figure 5B) demonstrates an ∼25% gain centrally, 25-50% gain through the intermediate regions, and ∼65% gain in the peripherally when comparing the proposed 16Tx/80Rx_2_ to our previous work, 16Tx/80Rx_1_.

**Figure 5:**
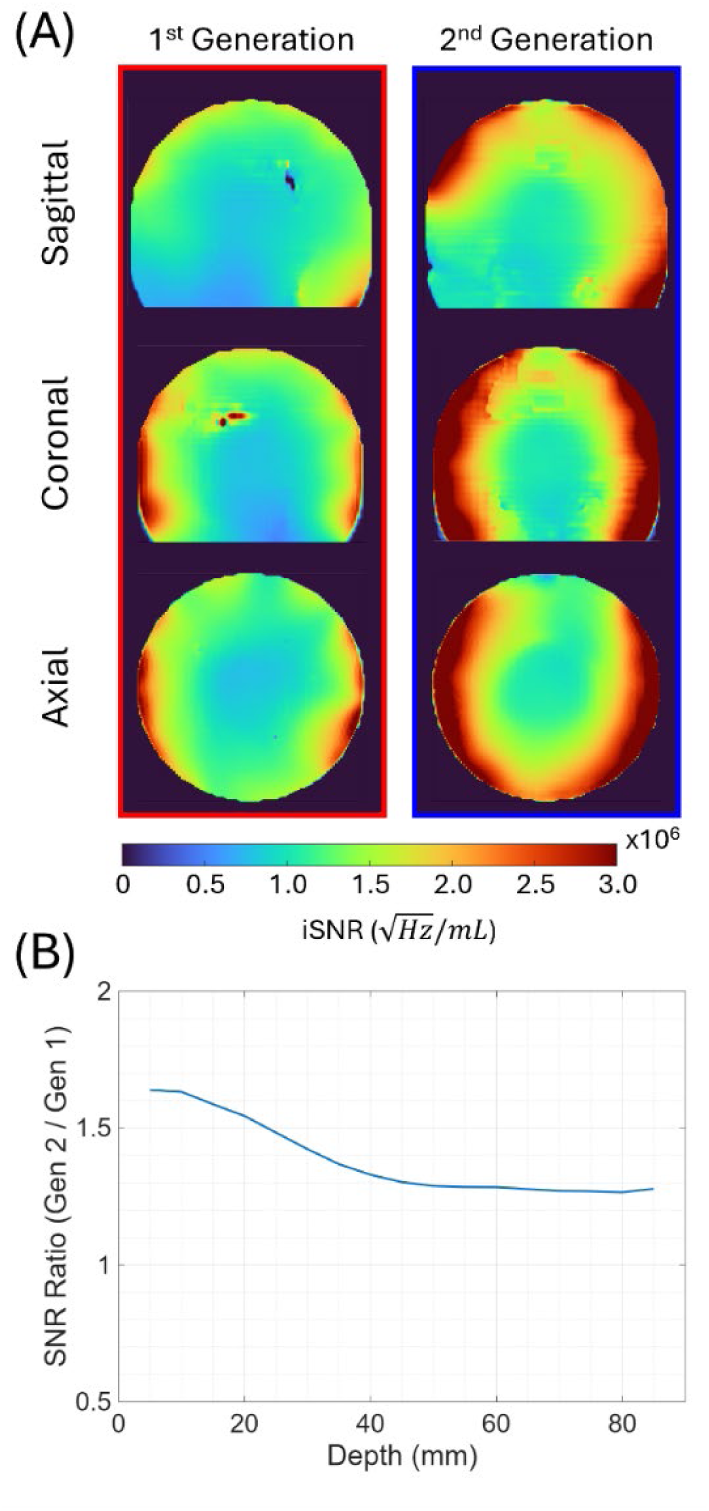
(A) SNR maps within the lightbulb-shaped phantom produced by the 1^st^ generation 16Tx/80Rx_1_ (Left) and 2^nd^ generation 16Tx/80Rx_2_ (Right) arrays. (B) The ratio of SNR as a function of depth plotted for SNR averaged in 0.5-cm-thick concentric shells and a 1-cm-diameter central volume. ranging from ∼25% centrally, 25-50% in the intermediate regions, and up to ∼65% peripherally.

Figure 6A shows SNR performance within the phantom in a central axial plane. The bar chart (Figure 6B) displays SNR performance as measured in a 2-cm-diameter central spherical volume. The updated 64Rx_2_ array captured 82% of uiSNR, while the previous generation (64Rx_1_) array captured 53%. The additional sensitivity from the 16-channel transceiver arrays increased those values to 91% and 71% respectively. For reference, data from the CMRR-built 7T 64-channel and 10.5T 128-channel receive arrays have been provided. (Figure S6B, S6D) shows the SNR performance plotted as function of depth for both the first- and second-generation 64Rx and 80Rx arrays.

**Figure 6:**
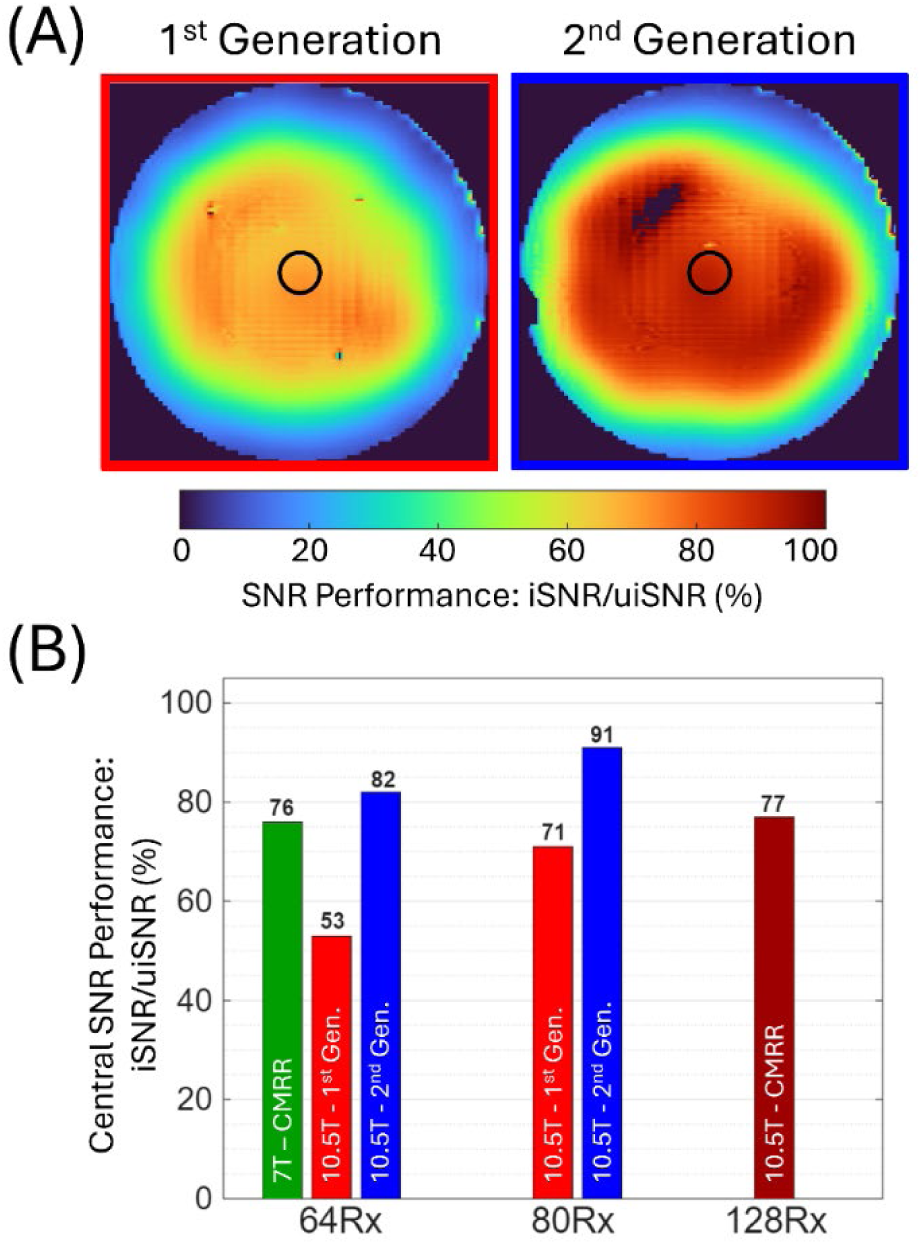
(A) The SNR performance of the arrays (left: 1^st^ generation 10.5T 16Tx/80Rx_1_, right: 2^nd^ generation 10.5T 16Tx/80Rx_2_) as a ratio of the ultimate-intrinsic SNR (uiSNR), presented as a map in a central axial slice of the lightbulb-shaped phantom. (B) Central SNR performance as measured in a 2-cm-diameter central spherical volume, identified by a black circle in (A). The uiSNR is different for 7T and 10.5T and was calculated separately for the 7T CMRR 64Rx (green), original 10.5T 64-, 80- (red), and 128Rx (maroon), and updated 10.5T 64- and 80Rx (blue) arrays at the two field strengths.

#### 3.2.3 In vivo SNR

Global SNR gains can be observed in vivo in Figure 7, which shows axial slices through the entire head. Here, we perceive similar central, intermediate, and peripheral SNR gains as observed in the phantom.

**Figure 7:**
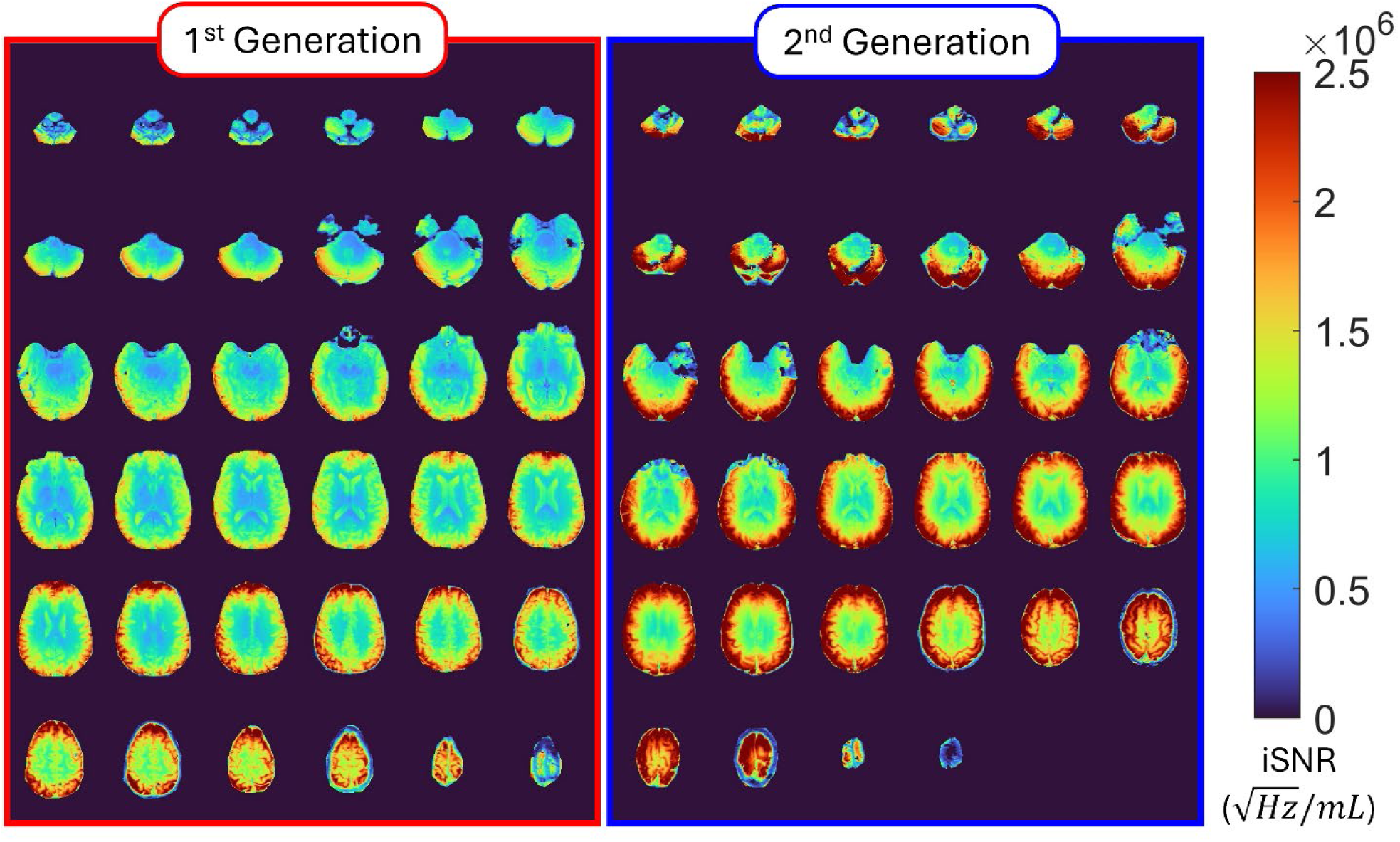
10.5T in vivo SNR gains achieved using the 2^nd^ generation 16TX/80Rx_2_ (Right) vs. the 1^st^ generation 16Tx/80Rx_1_ (Left) assembly. SNR data from both 16Tx/80Rx arrays are shown for two different human subjects.

#### 3.2.4 g-Factor and parallel imaging

The 64 receive-only channels are extracted from the 80-channel acquisition. To extract the receivers only, the signals from all channels are first decorrelated, and then the channels labelled as receivers are extracted. The 1/g factor is calculated using sensitivity profiles and the g-factor equation derived for a SENSE reconstruction. Figure 8 shows the 1/g maps over an 80mm central slab for the first- and second-generation arrays using a MIP through the 80mm slab.

**Figure 8:**
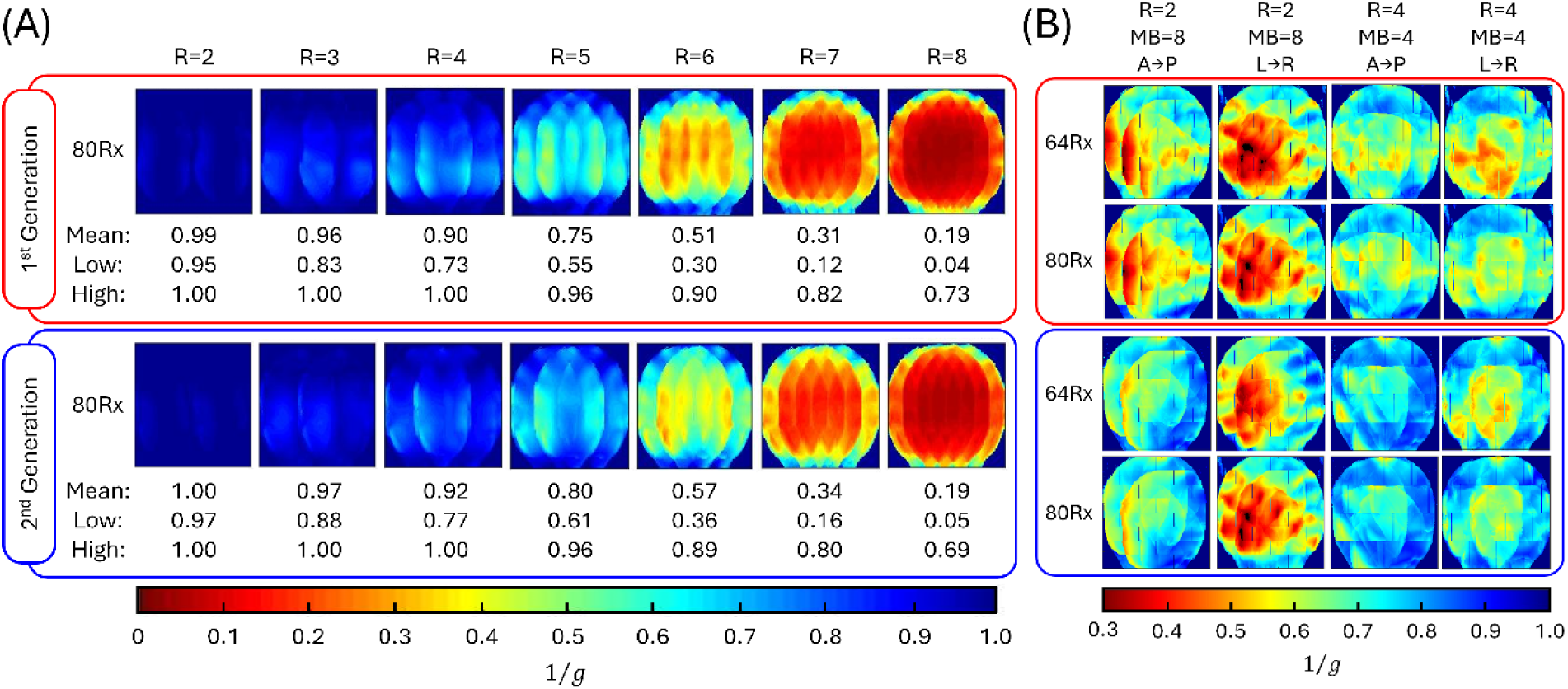
(A) Measured one-dimensional reduction factors (1/g maps) in the sagittal plane with the reduction factor is in the anterior-to-posterior direction for both 16Tx/80Rx array configurations. The low and high values listed are the 98^th^ percentile. (B) Experimental 1/g maps (sagittal & coronal orientations) over an 80mm central slab for different channel configurations and undersampling strategies using a MIP through the 80mm slab. Simultaneous multislice (SMS)/multiband (MB) acquisitions for MB= 8 & 4 in the superior-inferior (SI) direction and in-plane reduction factors of 2 and 4 in either the anterior-posterior (A→P) or left-right (L→R) direction.

Figure 8A presents 1-dimensional reduction performance, with the new 16Tx/80Rx_2_ array providing ∼10% gains in the mean 1/g values compared to the 16Tx/80Rx_1_ array with reduction factors of 5 through 8. Subtle gains can also be observed with the 16Tx/80Rx_2_ array when using 2-dimensional acceleration (Figure 8B).

#### 3.2.5 SNR across field strengths

Experimentally measured SNR maps in the lightbulb-shaped phantom for the four different array configurations, corrected for different T * at the two field strengths, are shown in Figure 9A. At 7T, relative to the industry-standard 32-channel array, the state of-the-art 64-channel array demonstrated a 47% increase in peripheral SNR, which tapered to an ∼4% increase in the center of the phantom. At 10.5T, the first-generation 80-channel array (16Tx/80Rx_1_) produced a 64% increase in peripheral SNR, and a 2-fold improvement centrally; relative the 7T standard, while the second-generation 80-channel array (16Tx/80Rx_2_) produced a 2.7-fold SNR increase in the periphery and 2.54-fold SNR increase in the central volume (Figure 9B).

**Figure 9:**
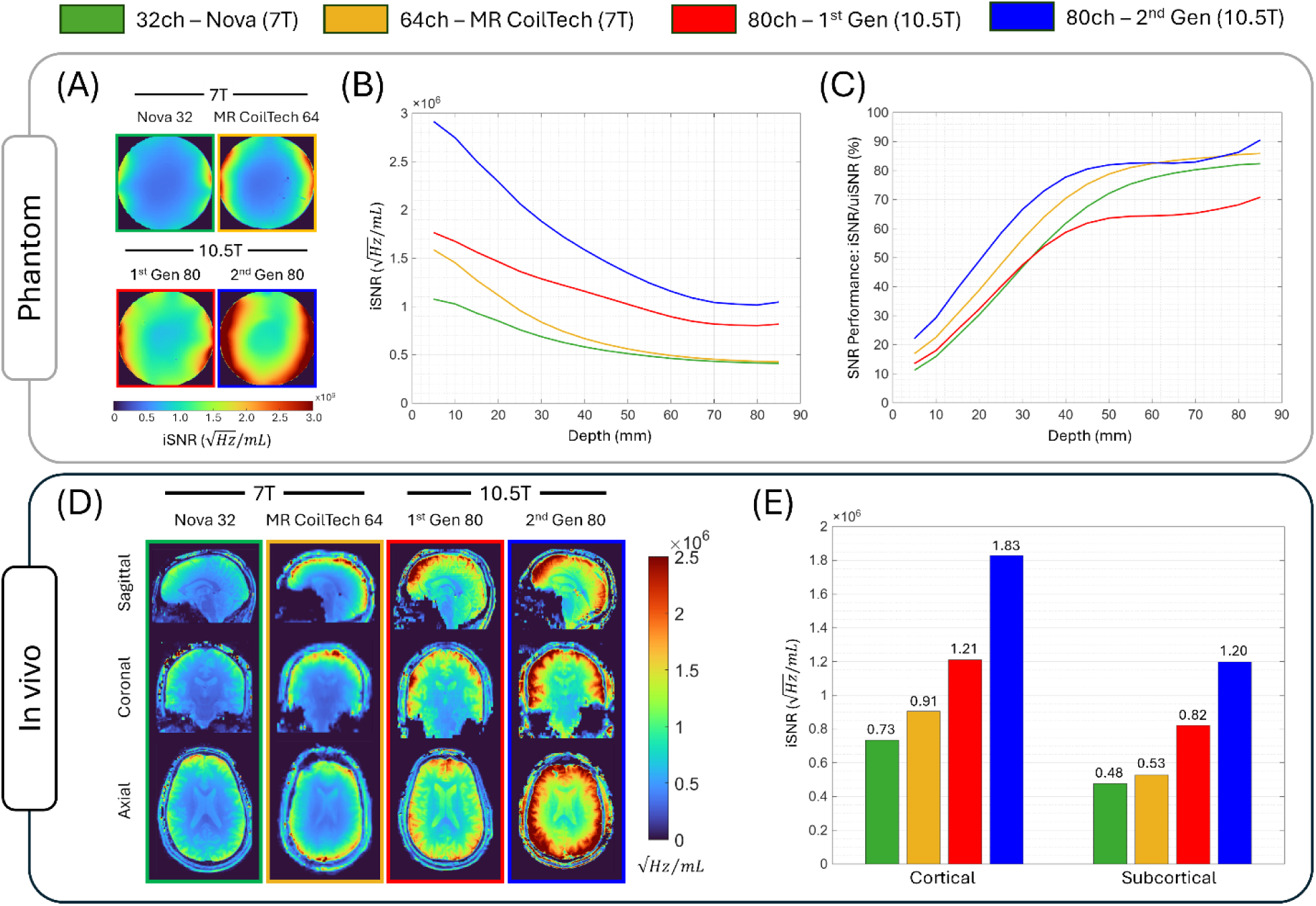
Intrinsic SNR (iSNR) and SNR performance (defined as iSNR/uiSNR) comparison across 7T & 10.5T field strengths and various RF coils. (Top Panel) phantom SNR maps of a central axial slice within the lightbulb-shaped phantom (A), as well as a function of depth plotted for SNR averaged in 0.5-cm-thick concentric shells and a 1-cm-diameter central volume (B). (C) Shows SNR performance as percentage of uiSNR within the same concentrical shells. (Bottom Panel) In vivo SNR maps from different subjects in all three orthogonal planes (D). In vivo SNR was quantified by splitting the brain into two large regions (cortical, and subcortical), and measuring the average SNR within those volumes (E).

In vivo SNR maps measured in different subjects (Figure 9D) show global SNR gains throughout the human head, including the brain stem. Intrinsic SNR (Figure 9E) is displayed when measured across the cortical and subcortical regions, demonstrating similar results as observed within the phantom.

SNR performance within the lightbulb-shaped phantom of the four arrays is shown in Figure 9C, with the 7T standard 32-channel array acquires 82%, the 7T state-of-the-art 64-channel array demonstrates 86%, the 10.5T first-generation 80-channel array obtains 71%, and the second-generation 80-channel array achieving over 90% of uiSNR in the central volume.

### 3.3 Neuroimaging at 10.5T

#### 3.3.1 Anatomical imaging

Figure 10 displays some in vivo utility of the new 16Tx80Rx_2_ array. Whole-head coverage is demonstrated with the sagittal MP2RAGE were acquired in ∼3 min, which requires a robust coil to handle the demanding 180 pulses and to generate spatially uniform contrast across the head within SAR limitations.

**Figure 10:**
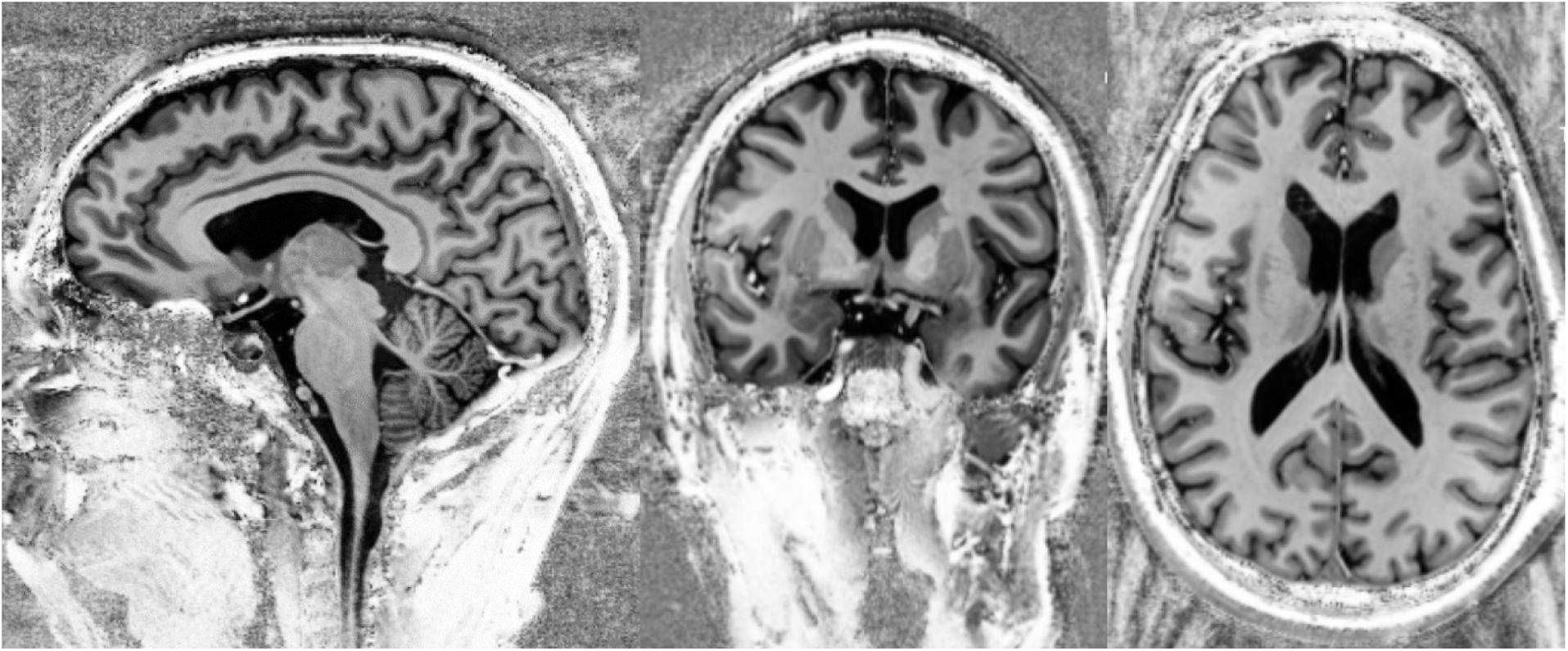
Examples of anatomical images at 10.5T obtained with the new 16Tx/80Rx_2_ array. Sagittal MP2RAGE image demonstrating whole-head coverage acquired in ∼3 mins.

Cerebellar-focused GRE images (Figure 11) with 0.2 × 0.2 × 1.0 mm^3^ resolution were acquired in ∼5 min. All images were acquired in the parallel-transmission mode using phase and amplitude B_1_ shim

**Figure 11:**
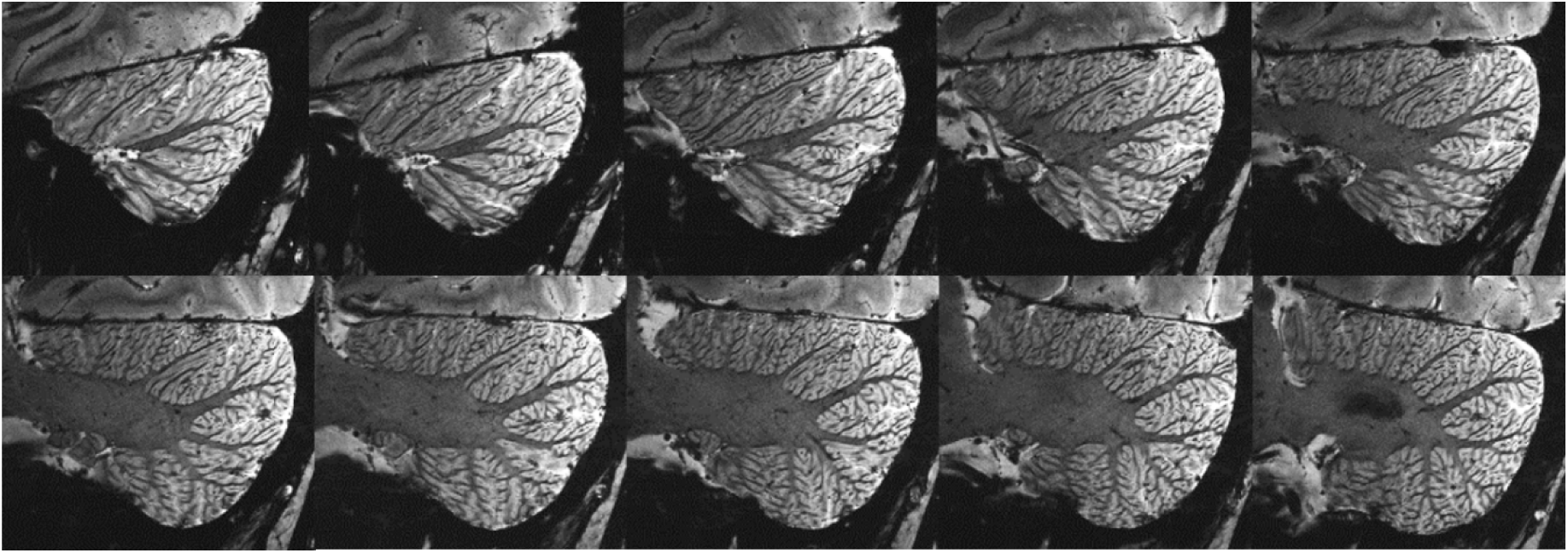
Cerebellar-focused sagittal 2D GRE with 0.2 × 0.2 mm^2^ in plane resolution.

Figure 12 displays SWI at 10.5T using a 3D-GRE sequence, and the minimum intensity projection for a 10.4-mm-thick slab from the data, acquired in ∼7 min using the new 16Tx/80Rx_2_ head array.

**Figure 12:**
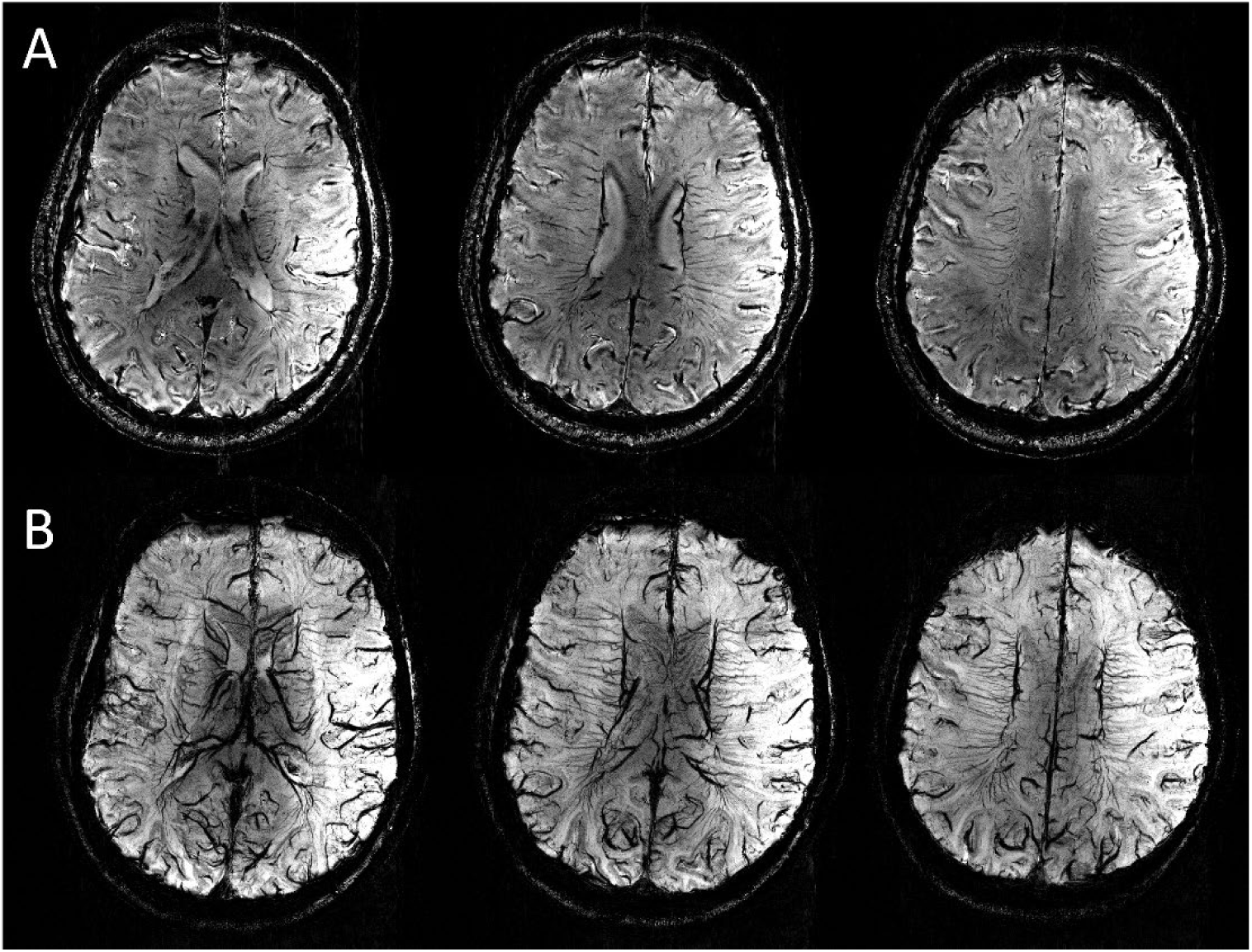
(A) Three slices from susceptibility weighted imaging (SWI) acquired in 36 contiguous, axial-coronal oblique slices with 0.21 × 0.21 mm^2^ in-plane and 1.3-mm slice resolution using a 3D-GRE sequence. (B) Minimum intensity projection (SWI) for a 10.4-mm-thick slab from these GRE data.

#### 3.3.2 Functional imaging

Figure 13 Demonstrates 3-plane BOLD EPI images with 0.6 mm isotropic resolution (0.216µL voxel volume) acquired using a whole-brain phase and amplitude B_1_ shim.

**Figure 13:**
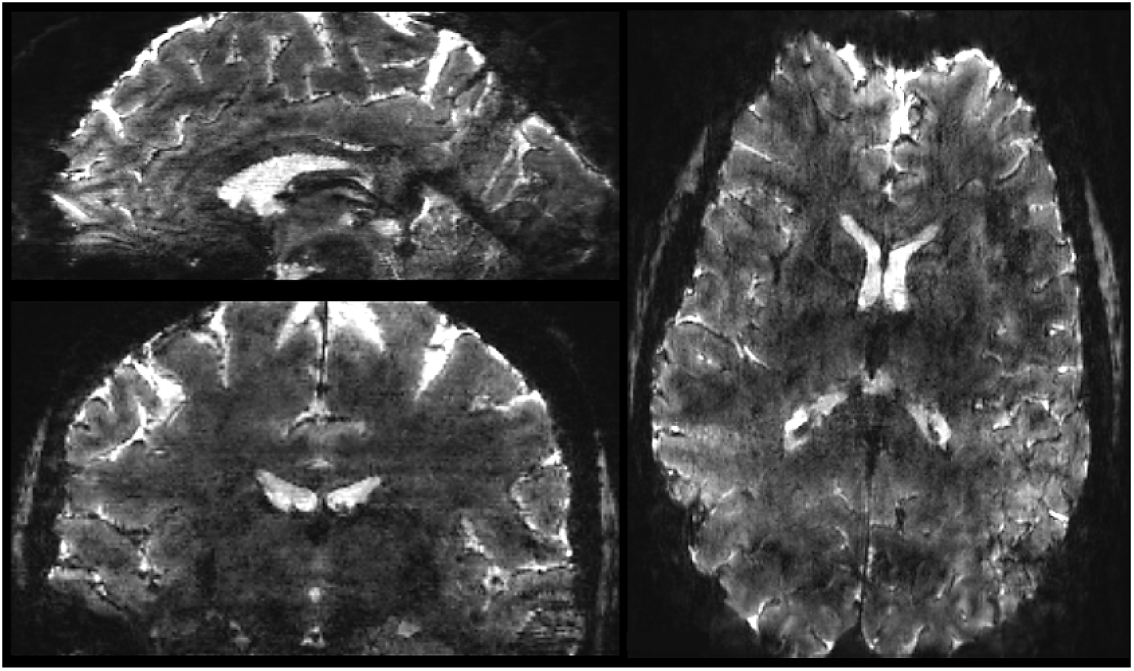
2D EPI images using 0.6 mm isotropic resolution, shown in all three orthogonal planes: sagittal (top-left), coronal (bottom-left),), and axial (right).

Figure 14 Demonstrates a mosaic of BOLD EPI images with 0.85 mm isotropic resolution acquired using a whole-brain phase and amplitude B_1_ shim.

**Figure 14:**
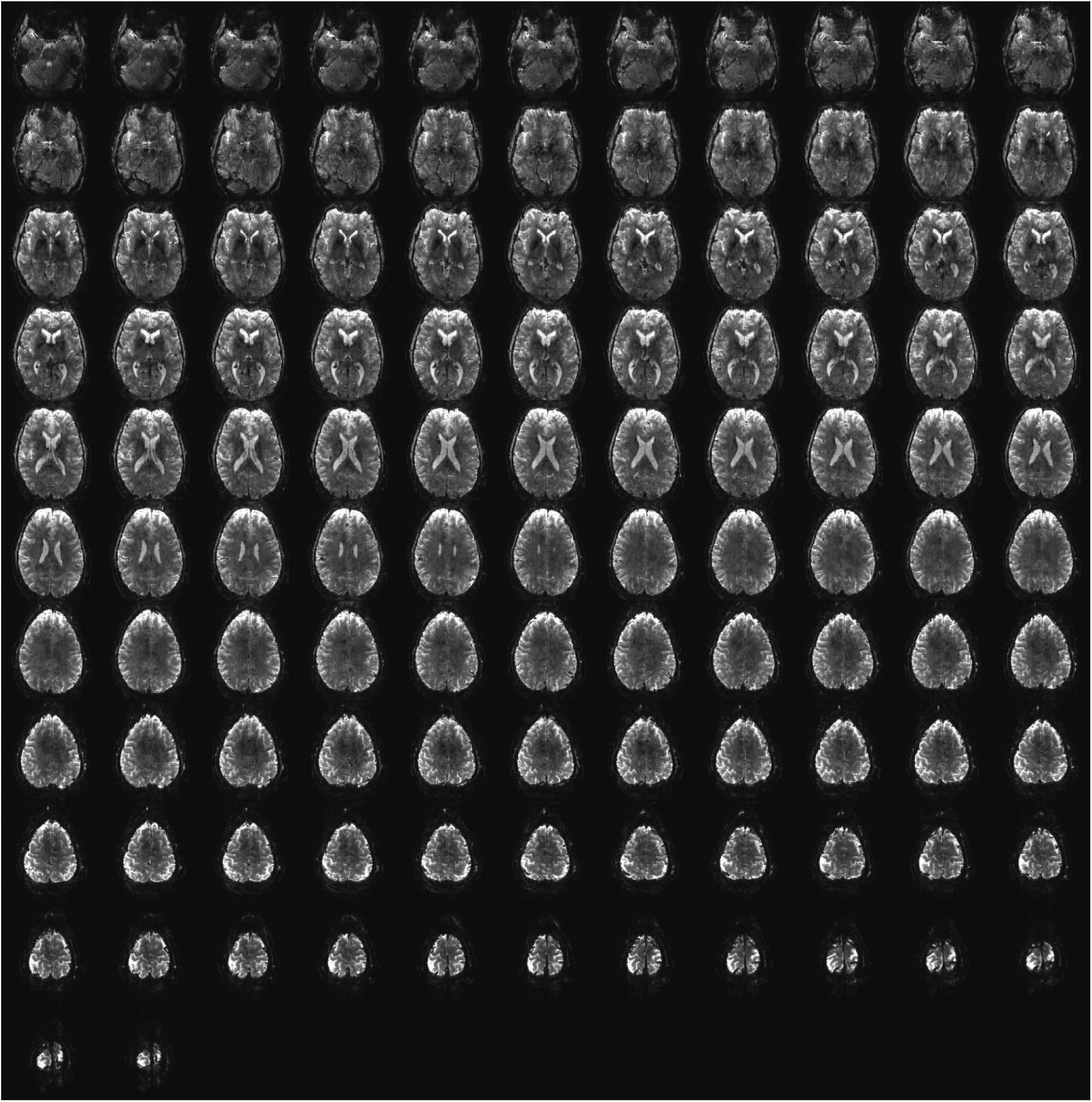
2D BOLD EPI images using 0.85 mm isotropic resolution.

## 4 Discussion

### Coil loss investigation

We have shown that careful design consideration should be given not only to receive coil size and array layout, but also to the parasitic losses that may accumulate, and become impactful as the receive coils inevitably move further from the sample due to various head sizes and shapes (Figure 4). The largest of those parasitic losses in the original circuit design was attributed to variable capacitors, which were used for tuning, and included in the distributed capacitor contribution. In our updated design, those variable capacitors were replaced with fixed ceramic-chip capacitors, which have lower losses. In this case, one is trading convenience for performance. Our solutions to minimize each of the parasitic losses resulted in improved Q-factors, which led to the design of the second-generation 64Rx_2_ array (Figure 2). These decreases in parasitic losses, combined with the increased filling factor of the larger Rx coil elements of the second-generation array led to an increase in Q_r_, which ultimately resulted in SNR improvements over the first-generation array of ∼25% centrally, 25-50% in the intermediate regions, and up to 65% peripherally within a phantom (Figure 5); similar gains were observed in vivo (Figure 7).

### SNR performance

In the first-generation of high-channel-count 10.5T coils (i.e., the 80-channel and 128-channel arrays), the realized central SNR performance of the receive-only elements (i.e., the 64-channel and 112-channel arrays) tiled over a head-conformal inner former was substantially lower than numerically predicted. Specifically, Vaidya et al. predicted that a 32-channel loop coil could capture more than 90% of the central uiSNR from 3T through 9.4T. In contrast, the first-generation 64- and 112-channel 10.5T loop arrays captured only 53% and 66% of the central uiSNR at 10.5T, respectively. To partially address this limitation, the first-generation 10.5T coils incorporated 16 relatively large self-decoupled loops as transceivers. Because of their nonuniform current distributions, these elements exhibit both dipole-like and loop-like behavior. This simple strategy recovered a substantial portion of the lost central SNR—which is now understood to have been limited by parasitic losses—and increased the central SNR performance of these arrays to 71% and 77%, respectively.

We continue to employ a 16-channel transceiver array in combination with this second-generation fully-overlapped high-channel-count receive array (Figure 3D) in order to increase SNR throughout the sample. Here we utilized an aligned LFD transceiver element design vs. the staggered SD element from our prior work. In this case, the SNR contribution of the transceiver coil was modest but still measurable. Specifically, the central SNR performance of the receive-only coil increased from 82% to 91% after addition of the transceiver array (Supporting Figure S6).

In addition to central SNR performance, Vaidya et al. predicted in the same numerical study that a 32-channel loop coil would capture more than 80% of the uiSNR in intermediate regions of the head up to 9.4T. However, the first-generation 10.5T coils (i.e., the 80-channel and 128-channel arrays) realized only ∼50% of the uiSNR in those regions. With the design improvements introduced in the second-generation 10.5T 80-channel coil, the realized performance in intermediate regions now reaches ∼80% of uiSNR. In summary, the second-generation 10.5T 80-channel coil now meets the numerical predictions of SNR performance in both central (∼90%) and intermediate (∼80%) regions (Figure 9C).

Nevertheless, SNR performance in peripheral regions where the uiSNR is intrinsically very high, remains relatively low (∼20%). This is a shortcoming that is shared by all arrays in general,^44^ including the 7T and 10.5T arrays we have examined in this work (Figure 9), though the second-generation 10.5T array has so far the best SNR performance at 20%. Thus, despite a significant improvement over prior work both at 7 and 10.5T, additional targeted strategies will be required to increase the SNR performance peripherally. These may include further increasing the number of loops while reducing the sample-to-coil distance through flexible coil technology, while still satisfying the Q-ratio considerations emphasized throughout this manuscript.

### Parallel imaging performance

Afore discussed peripheral to central SNR performance improvements in the second-generation CMRR 16Tx/80Rx_2_ coil, relative to the first-generation version, reflect only the intrinsic SNR advantage of the design under fully sampled k-space conditions. However, one of the primary motivations for developing such high-channel-count coils is to accelerate imaging through k-space undersampling, which inevitably amplifies noise and reduces image SNR. This noise amplification is commonly quantified as the g-factor, which depends on the geometry of the underlying Rx coil and, as a rule of thumb, often improves with increasing channel count when sensitivity encoding is enhanced. Other major factors that influence the g-factor are the degree of similarity between channel-sensitivities and phase patterns, as well as the noise correlation between channels: the more distinct the sensitivity and phase profiles of each individual coil and the lower the noise correlation, the better the g-factor performance. To achieve this, gaps have traditionally been introduced between neighboring loops, often in the circumferential direction, which is the strategy we had also taken in our first-generation 16Tx/80Rx_1_ array. This practice naturally reduces loop size. By contrast, the second-generation CMRR 80-channel coil used larger loops with overlap in all directions, which might have been expected to degrade g-factor performance. However, our findings in Figure 8 show the opposite: g-factors were not degraded and were even improved in some cases by approximately 10%.

This somewhat counterintuitive behavior can be understood by separating the contributing factors. First, noise correlation is related to electric coupling between loops, and as described by Roemer et al.,^22^ such electric coupling can only be reduced by increasing the separation between elements. From this perspective, the tighter and more overlapped geometry of the second-generation 80-channel coil would be expected to work against g-factor improvement. Second, shared sensitivity can arise for two reasons. One is purely geometric overlap in the right-handed rotating magnetic fields (B_1_^−^) of neighboring loops. Because the phase of the B_1_^−^ fields vary more rapidly in space at higher field strengths, this effect is generally less severe at 10.5T than at lower fields, and thus does not strongly favor either generation of coils. The second, and more important, source of shared sensitivity is induced current generated by magnetic coupling between loops. Magnetic coupling can be minimized to nearly zero either by precise geometric overlapping of adjacent loops or, in non-overlapping layouts, through preamplifier decoupling. However, the effectiveness of preamplifier decoupling depends not only on accurate implementation but also strongly on the initial level of coupling between elements, as described by Roemer et al.^22^ In the case of non-overlapping but adjacent loops, the starting level of coupling can substantially compromise the effectiveness of preamplifier decoupling, leading to induced currents in neighboring loops and, consequently, increased similarity in their sensitivity profiles. This mechanism likely represents a major contributor to the improved g-factor performance of the second-generation coil, in which, both geometric and preamplifier decoupling technique were carefully implemented.

In summary, in the parallel imaging regime, the new second-generation coil provides not only a 25%–65% intrinsic SNR gain over the first-generation coil, but also an additional ∼10% improvement associated with more favorable parallel imaging performance.

### Field dependence of SNR

The inherently higher available SNR (i.e. higher uiSNR) at higher field strengths has been one of the primary motivations for using UHF MRI scanners. It is therefore valuable to understand exactly how much of these SNR gain become available as B_0_ increases. Although the literature contains a wide range of experimentally measured SNR gains with B_0_, there is general consensus that the SNR increase is supralinear beyond 3 Tesla, as supported by uiSNR theory and numerical calculations.^1,5,67^ The dependence of uiSNR on B_0_ varies with object type, object geometry, and region of interest; however, for the human brain, it can generally be stated that the uiSNR dependence on B_0_ is close to linear near the periphery and increases to approximately quadratic toward the center. In fact, Pfrommer et al.^67^ showed in a numerical study spanning 3T to 11.7T with a realistic human head model that the threshold for linear B_0_-dependence of uiSNR lies approximately 16–20 mm beneath the skin and follows the contour of the skull. They further showed that, within the cerebrum, uiSNR increases supralinearly, with an exponent of 1.7 at an intermediate depth and 2.13 at the center. These findings are consistent with our previous numerical study at 7T and 10.5T by Zhang et al.,^1^ in which we observed B_0_-dependence exponents of 1.7 and 2.2 in intermediate and central brain regions, respectively. Le Ster et al.^5^ investigated the central SNR gain from 3T to 11.7T experimentally and using analytical calculations of uiSNR in a spherical phantom and reported an exponent of ∼2.

In the present study, the findings shown in Figure 9—comparing the second-generation 10.5T 16Tx/80Rx_2_ with a state-of-the-art commercially available 7T 64-channel coil—show that, at the center of a lightbulb-shaped phantom, the SNR gain follows a B_0_ exponent of 2.2. Similarly, for in vivo SNR measurements, exponents of 1.73 and 2.02 were calculated in cortical and subcortical regions, respectively, for these coils in close agreement with the numerical predictions reported by Pfrommer et al.,^67^ as well as in our previous work by Zhang et al.^1^ It is important to note that the B_0_-dependence of SNR reported here for 7T and 10.5T does not isolate the effect of field strength alone, as differences in Rx coil design—including channel count and layout—were not controlled for. Rather, this comparison reflects the performance of two state-of-the-art coils at their respective field strengths.

### Anatomical and functional imaging

Figures 10-14 demonstrate the utility of the new 16Tx/80Rx_2_ array coil for both anatomical and functional imaging. These preliminary results are not meant to explore the ultimate limits of resolution and/or contrast available at 10.5T; rather to illustrate the feasibility and potential that continual hardware developments may have on human brain imaging at UHF.

## 5 Conclusions

A new 16-channel transmit, 80-channel receive head coil comprising a 16-channel loop-folded dipole transceiver array and 64-channel receive-only coil array has been developed for neuroimaging applications at 10.5 T. This device was the result of an engineering redesign and optimization effort motivated by our previous works, and the results show significant improvements in transmit efficiency, parallel imaging, SNR, and SNR performance. We present solutions to many of the contributing factors of receive coil loss, which degrade performance, and demonstrate the ability to approach uiSNR limits. With the large gains in SNR and improved g-factors compared to our previous work, the accelerated SNR of the updated 16Tx/80Rx_2_ shows great promise for improved neuroimaging at 10.5 T.

## Acknowledgments

This research was funded by NIH grants UM1 NS132207, P41 EB027061, R01 EB038654, R01 EB031765, and S10 RR029672.

## Supporting Information

### Coil Loss Investigation

#### Experimental Setup

The Q ratios for coils with different components and architectures were calculated using individual unloaded (Q_U_) and loaded (Q_L_) Q factor measurements (with Q ratio equal to Q_U_/Q_L_). Different coil sizes (3x5 cm^2^and 5x5 cm^2^), matching networks (L-match and lattice network), active detuning architectures, passive detuning locations, and capacitor types (fixed and variable) were explored.

To easily evaluate and compare the differences between these components/networks, Q ratios were measured in incremental steps beginning with only the loop and distributed capacitors, then adding just the matching network, and finally the detuning network to create a complete loop.

For a comprehensive comparison, four loops were made: a 3x5 cm^2^ loop with an L-match, a 3x5 cm^2^ loop with a lattice match, a 5x5 cm^2^ loop with an L-match, and a 5x5 cm^2^ loop with a lattice match. Due to the different matching network, passive detuning network location differences were only measured on the 5x5 cm^2^ loop with a lattice match. Capacitor type differences were measured on a 3x5 cm^2^ loop with just distributed capacitors, where one capacitor is swapped out for a variable capacitor.

For all unloaded measurements, all loops were placed above the center of a large copper sheet with a 5-cm thick polyethylene foam pad as a spacer, as shown in Figure S1 (left). The copper sheet and 5-cm foam spacer were then removed and loaded measurements were made with the loops placed on top of a cube-shaped phantom (σ = 0.97 S/m, ε_r_ = 80) at various distances ranging from 1-cm to 4-cm, with separate 1-cm foam spacers used for coil-phantom separation, as shown in Figure S1 (right).

All measurements were taken using a decoupled double probe pair, with the probe placed above the loops at a distance far enough away to avoid loading the loops, but close enough to measure a strong signal (found to be ∼4 cm when unloaded and ∼2 cm when loaded), in addition to the probe being secured by a vise to ensure placement above each loop’s center and consistency between measurements.

#### Results

Tables containing the Q ratio (Q_U_/Q_L_) values for coils with various sizes and matching/detuning networks are shown below. Note the increased Q ratio of the 5x5 cm^2^, lattice match loops compared to the 3x5 cm^2^, L-match loops, with the 5x5 cm^2^, lattice match loop seeing an average Q ratio increase across all distances of 48.9% when complete with matching and detuning networks, as shown in Table S1.

For coils with a lattice matching network, a passive detuning network is needed and to evaluate the best position, Q ratio measurements were taken with the network at the loop’s input (i.e. parallel to the PIN diode used in the active detuning network) and parallel to the distributed capacitor at the top of the loop. The 5x5 cm^2^, lattice match loop placed 1-cm from the phantom was used for this comparison, where the passive detuning network at the input showing a 26.3% higher Q ratio compared to the network at the top of the loop, as shown in Table S2.

Using a 3x5 cm^2^ loop with only distributed capacitors, the impact of the inclusion of a variable capacitor was explored, with the results shown in Table S3. By excluding the variable capacitor, the Q ratio over all distances saw an average increase of 42.1%.

---

**FIGURE S1:**
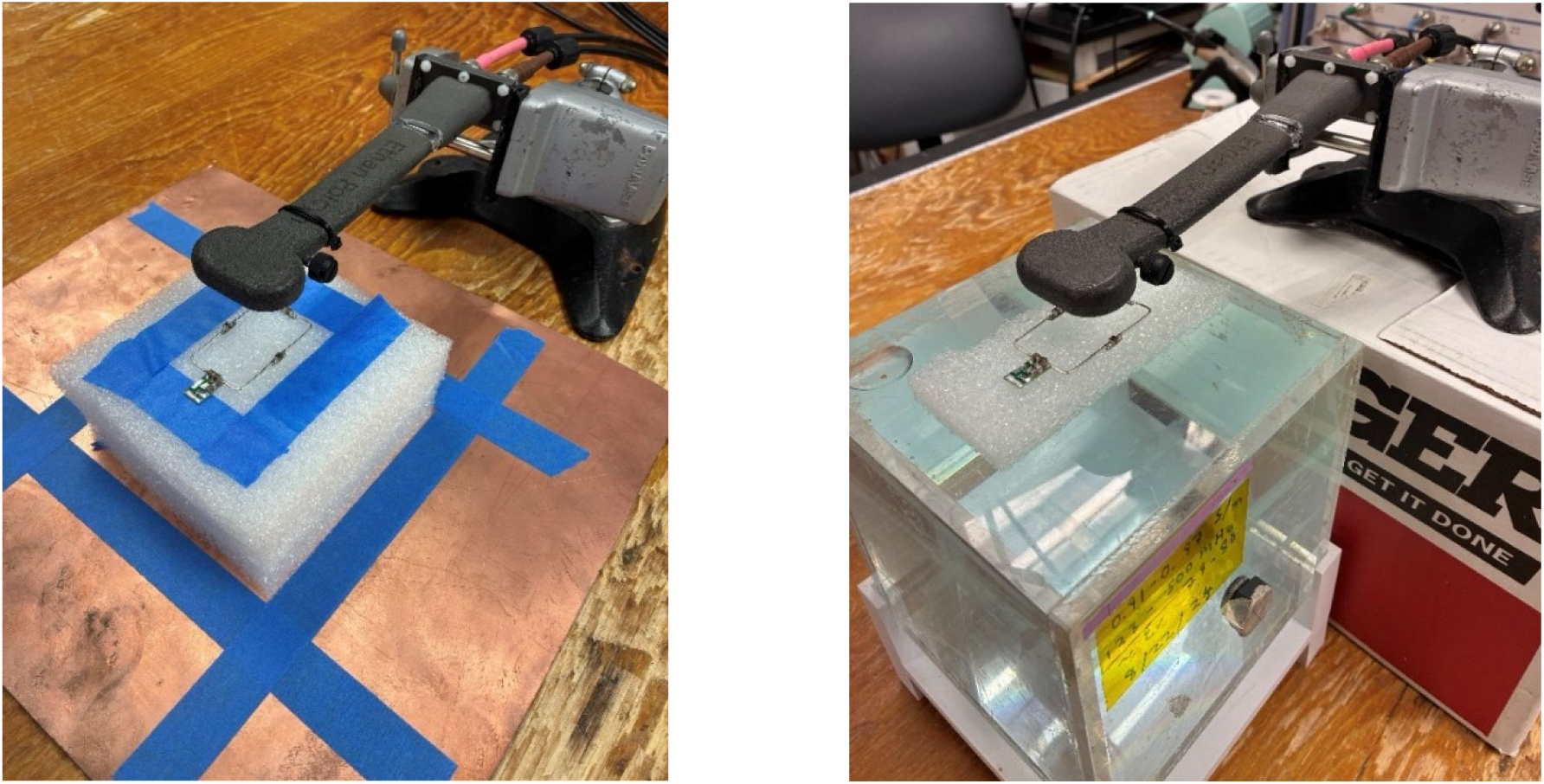
Images showing the experimental setup for unloaded (Left) and loaded (Right) Q measurements for a 3x5 cm, L-match loop with the loop and matching network and 1-cm coil-phantom distance.

**FIGURE S2:**
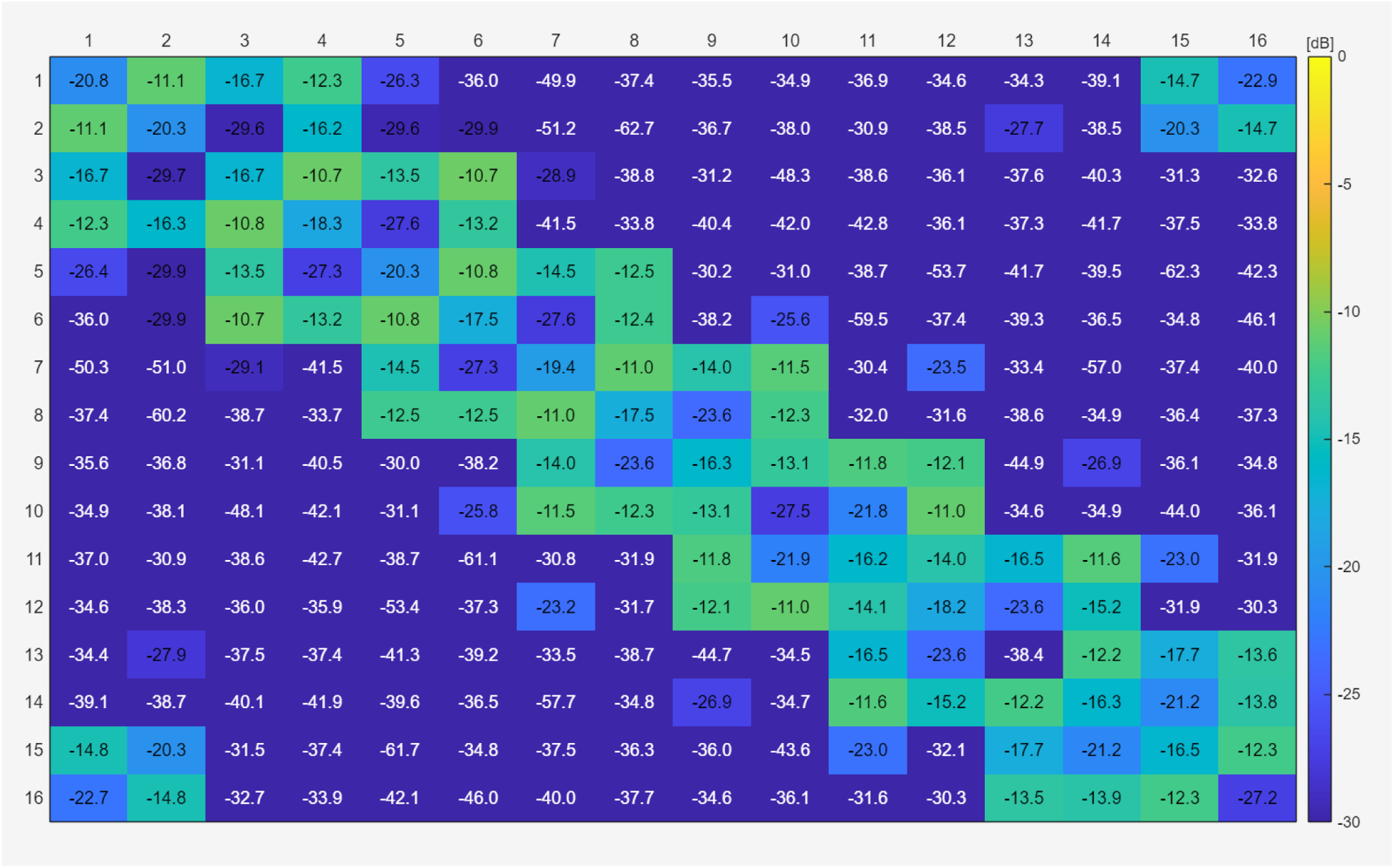
S-parameter matrix for the 16Tx/Rx_2_ LFD array with the updated 64Rx_2_ array while loaded with the lightbulb-shaped phantom.

**FIGURE S3:**
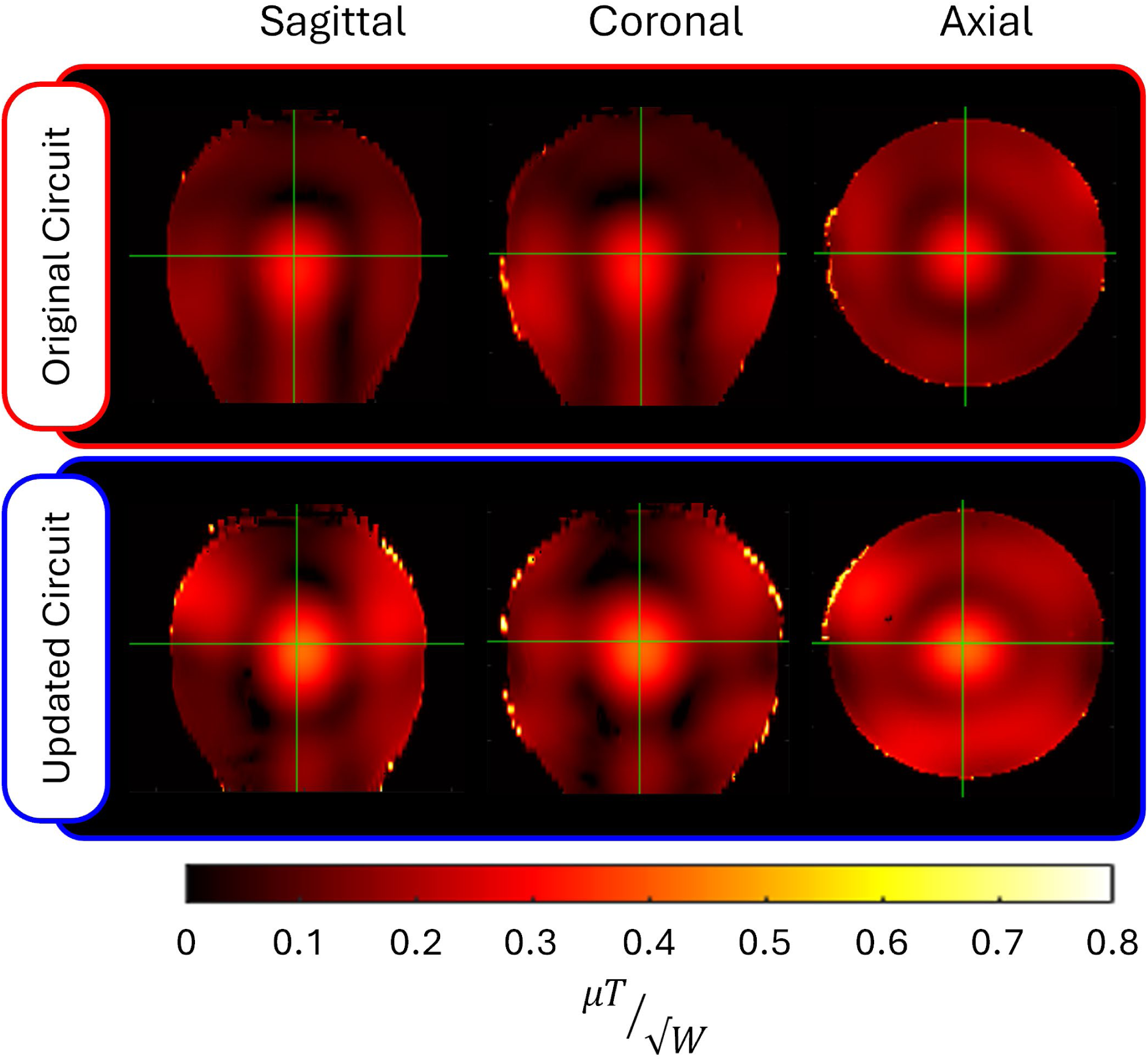
B_1_^+^ efficiency maps within the lightbulb-shaped phantom produced by the 1^st^ generation 16Tx/80Rx_1_ (Top) and 2^nd^ generation 16Tx/80Rx_2_ (Bottom) arrays, demonstrating 0.313 and 0.414 µT/√W, respectively when measured in a 1-cm diameter spherical volume centered at the crosshairs within the phantom.

**FIGURE S4:**
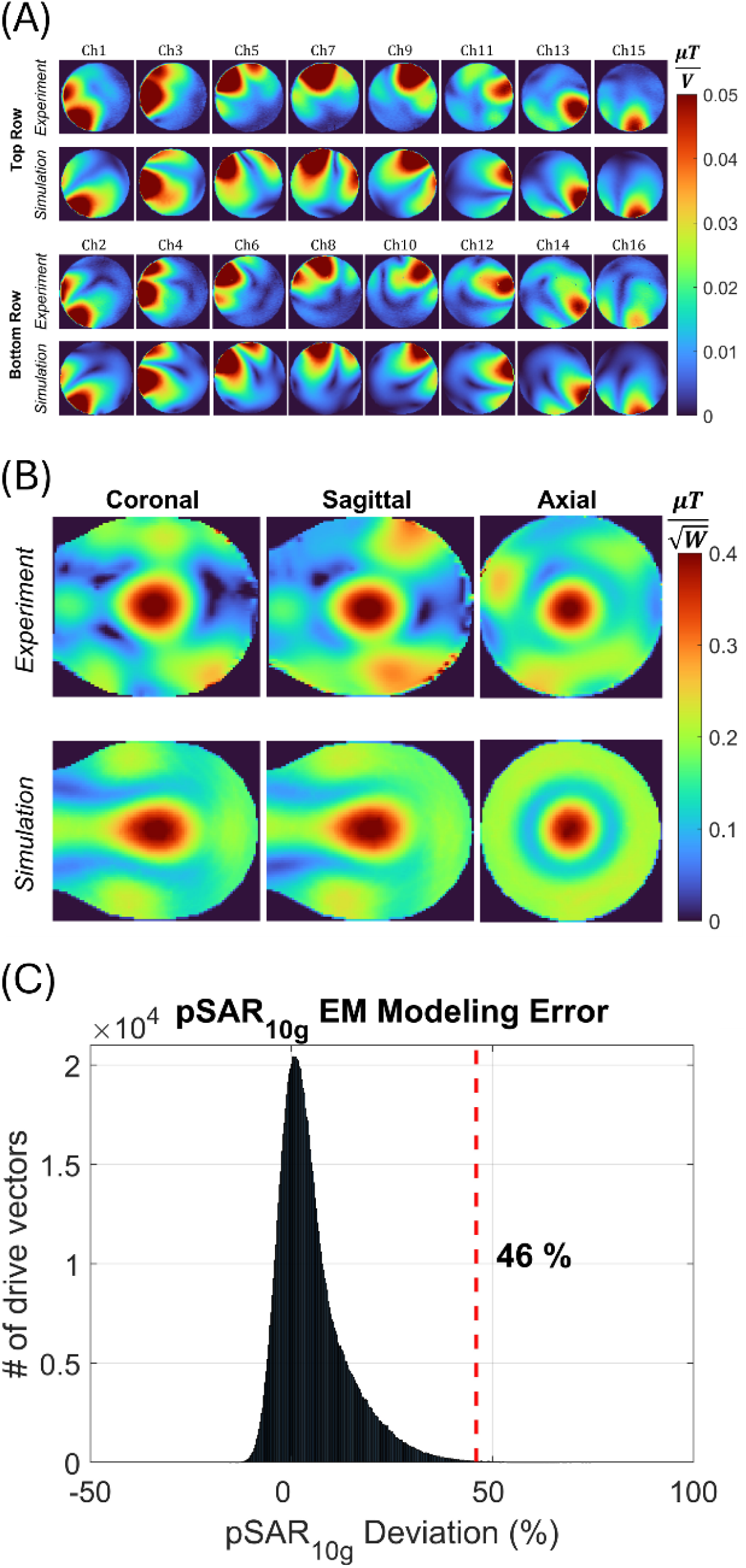
(A) Shows per-channel comparison of B1+ between simulation and experiment. These fields were used to generate the circularly polarized (CP)–like B_1_^+^ maps (B) for the complete assembly, a normalized root mean square error of 35% was calculated between simulation and experiment. (C) Shows the predicted error in peak 10 g-averaged local specific absorption rate. When combined with intersubject variability and power monitoring uncertainty, yielded a safety factor of 1.7.

**FIGURE S5:**
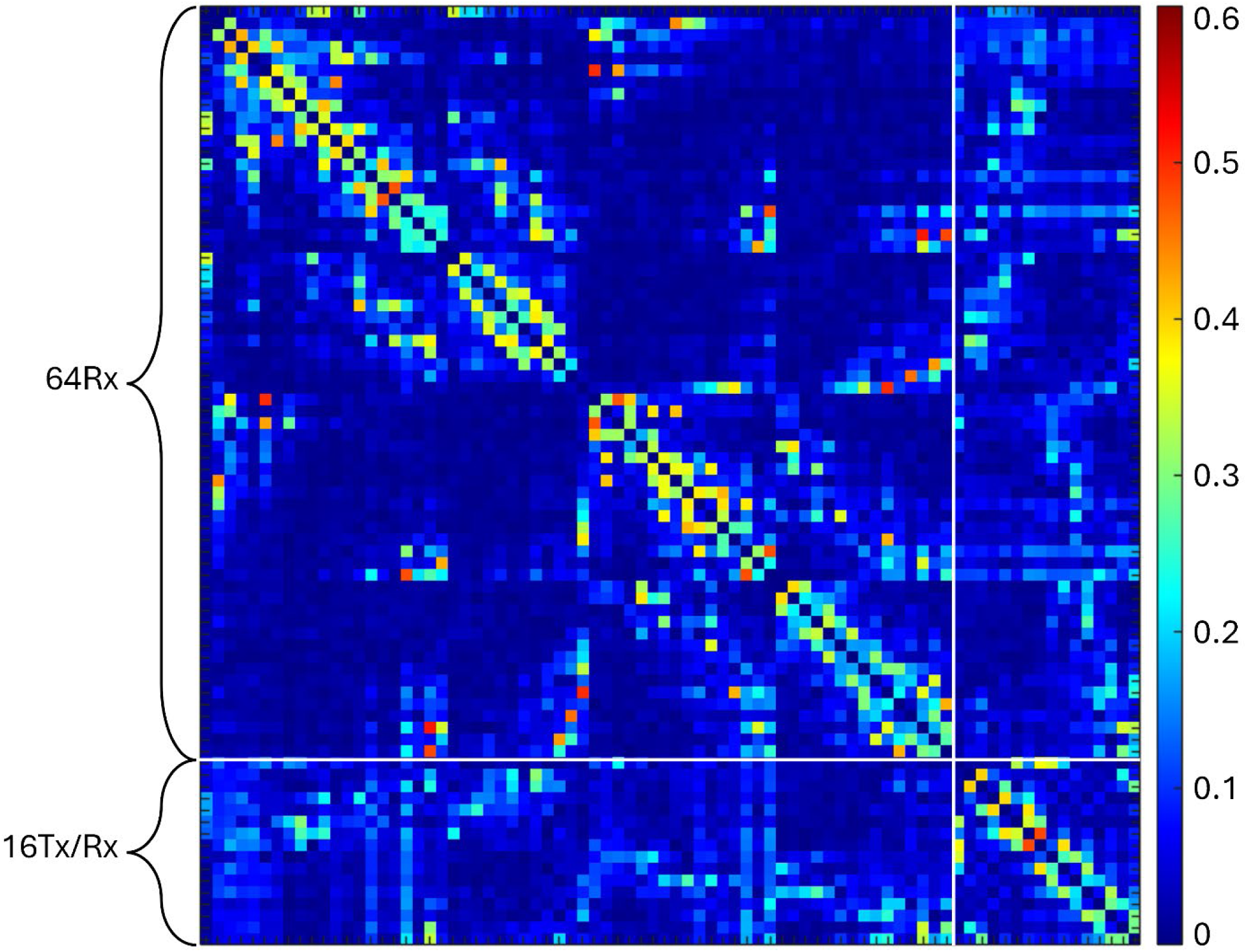
Experimentally measured noise correlation matrix of the 10.5T 16Tx/80Rx_2_ array while loaded with the lightbulb phantom (80Rx channels are composed of the 64-channel receive-only array paired with the 16-channel transceiver array). The average noise correlation for 80-channels was 6.1% with the maximum observed being 51%.

**FIGURE S6:**
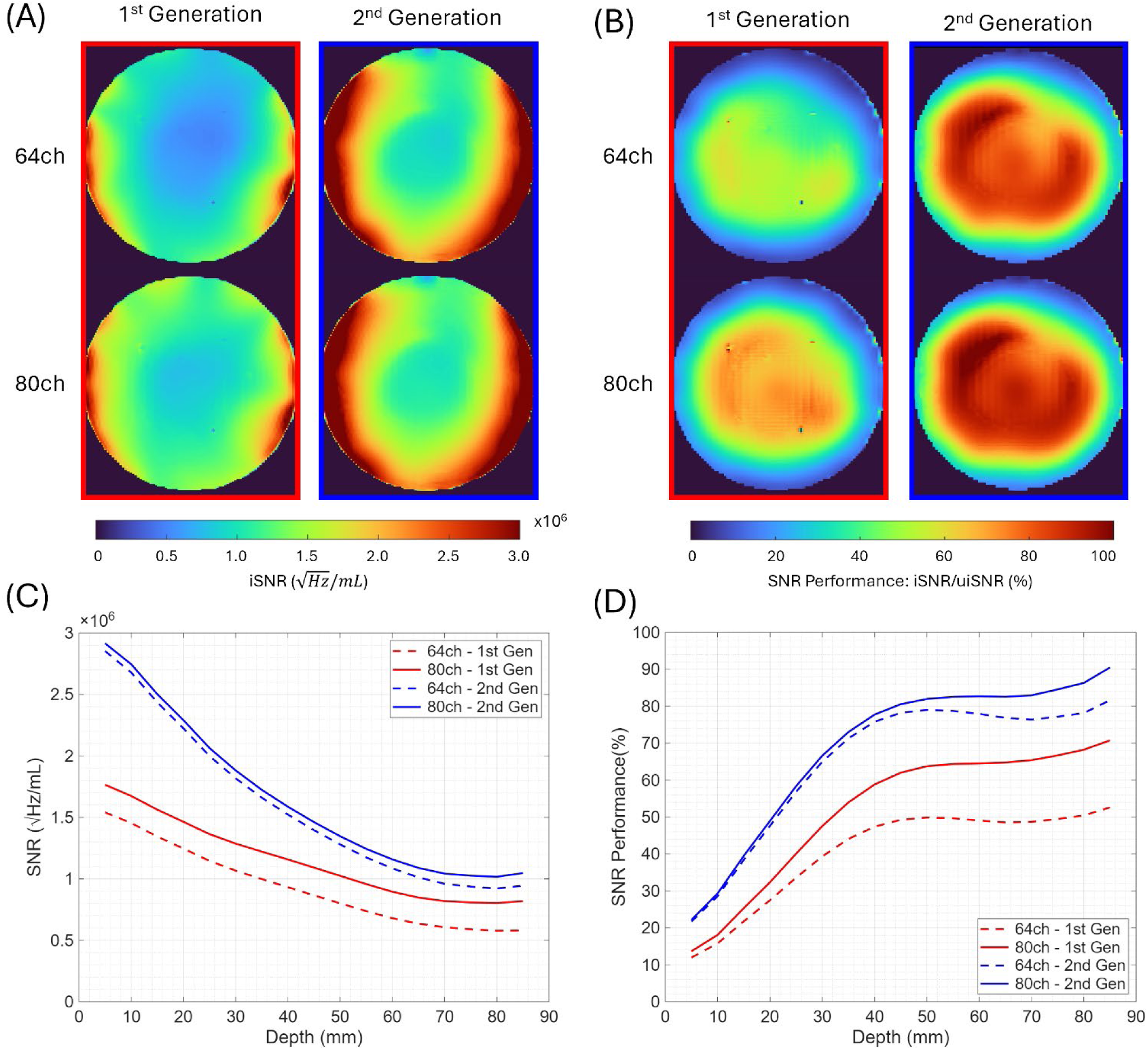
SNR comparison between the 1st and 2nd generation arrays. (A) SNR maps showing the contributions from the 64-channel receive only array (A, Top), and complete 80-channel assembly (A, Bottom) in the central axial slice of the lightbulb-shaped phantom. (C) shows the SNR as a function of depth plotted for SNR averaged in 0.5-cm-thick concentric shells and a 1-cm-diameter central volume. (B) Shows SNR performance as percentage of uiSNR within the same axial slice, (D) shows SNR as a function of depth within the same concentrical shells.

**Table S1:**
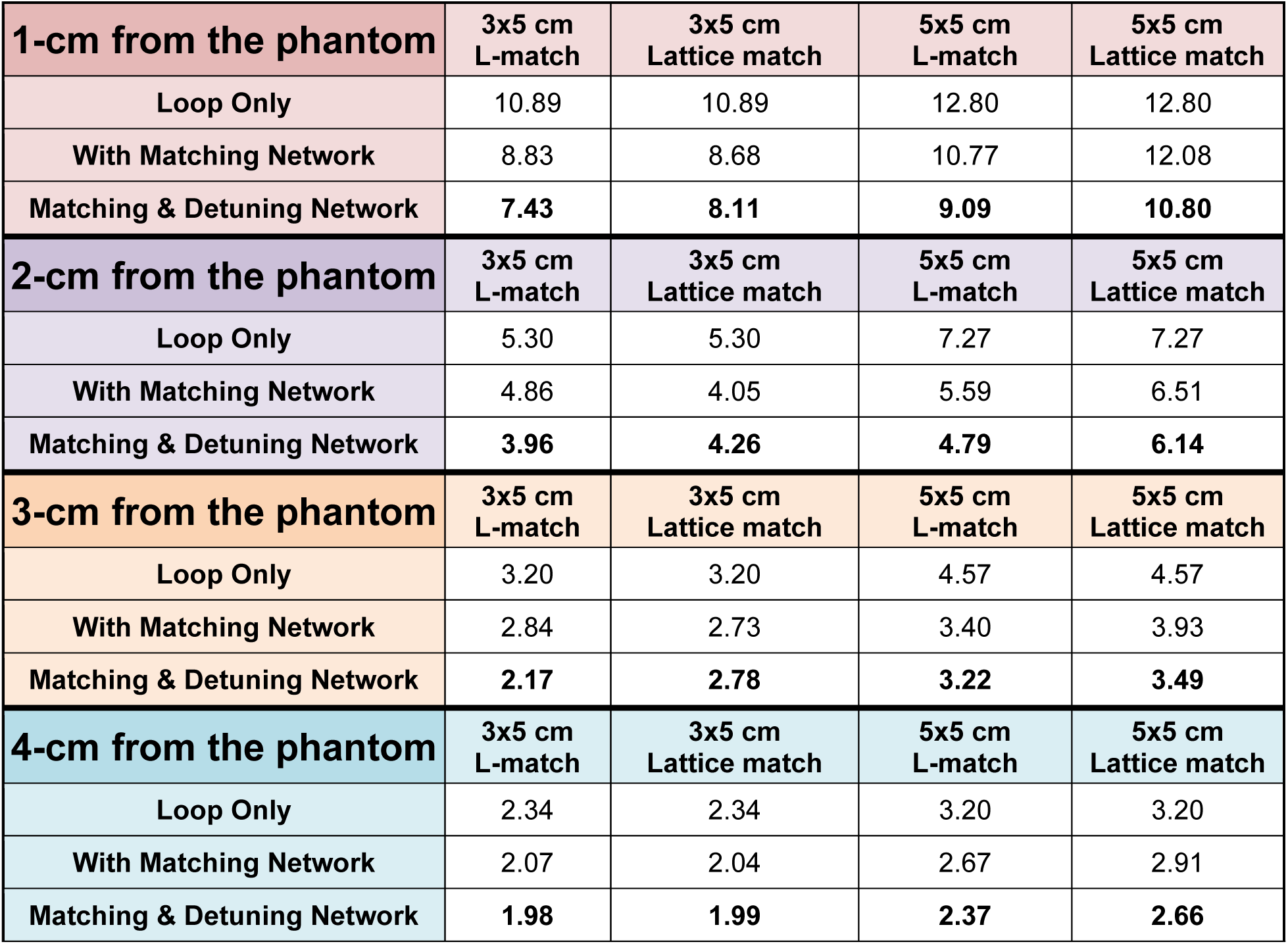
Q Ratio values with coils loaded 1 to 4 cm from the phantom for each loop type in incremental steps throughout coil building.

| **1-cm from the phantom** | **3x5 cm**<br>**L-match** | **3x5 cm**<br>**Lattice match** | **5x5 cm**<br>**L-match** | **5x5 cm**<br>**Lattice match** |
| --- | --- | --- | --- | --- |
| Loop Only | 10.89 | 10.89 | 12.80 | 12.80 |
| With Matching Network | 8.83 | 8.68 | 10.77 | 12.08 |
| Matching & Detuning Network | **7.43** | **8.11** | **9.09** | **10.80** |
| **2-cm from the phantom** | **3x5 cm**<br>**L-match** | **3x5 cm**<br>**Lattice match** | **5x5 cm**<br>**L-match** | **5x5 cm**<br>**Lattice match** |
| Loop Only | 5.30 | 5.30 | 7.27 | 7.27 |
| With Matching Network | 4.86 | 4.05 | 5.59 | 6.51 |
| Matching & Detuning Network | **3.96** | **4.26** | **4.79** | **6.14** |
| **3-cm from the phantom** | **3x5 cm**<br>**L-match** | **3x5 cm**<br>**Lattice match** | **5x5 cm**<br>**L-match** | **5x5 cm**<br>**Lattice match** |
| Loop Only | 3.20 | 3.20 | 4.57 | 4.57 |
| With Matching Network | 2.84 | 2.73 | 3.40 | 3.93 |
| Matching & Detuning Network | **2.17** | **2.78** | **3.22** | **3.49** |
| **4-cm from the phantom** | **3x5 cm**<br>**L-match** | **3x5 cm**<br>**Lattice match** | **5x5 cm**<br>**L-match** | **5x5 cm**<br>**Lattice match** |
| Loop Only | 2.34 | 2.34 | 3.20 | 3.20 |
| With Matching Network | 2.07 | 2.04 | 2.67 | 2.91 |
| Matching & Detuning Network | **1.98** | **1.99** | **2.37** | **2.66** |

**Table S2:** Q Ratio values for a 5x5 cm^2^, lattice match loop loaded 1-cm from the phantom with the passive detuning network placed at different points on the loop.

|  | No Passive Detuning | Passive Detuning at input (% Lost) | Passive Detuning at top of loop (% Lost) |
| --- | --- | --- | --- |
| **Q Ratio** | 10.75 | 9.52<br>(11.4%) | 7.54<br>(29.9%) |

**Table S3:** Q Ratio values for a 3x5 cm^2^ loop with only distributed capacitors at various loading conditions, where one of the distributed capacitors is changed from a fixed to variable capacitor.

| Coil-to-Phantom Distance | 3 Fixed, 1 Variable | All Fixed | % Improvement w/o Variable Capacitor |
| --- | --- | --- | --- |
| **1-cm** | 7.22 | 10.89 | 33.7 |
| **2-cm** | 3.61 | 5.30 | 31.9 |
| **3-cm** | 2.28 | 3.20 | 28.8 |
| **4-cm** | 1.79 | 2.34 | 23.5 |

